# Variable brain wiring through scalable and relative synapse formation in *Drosophila*

**DOI:** 10.1101/2021.05.12.443860

**Authors:** F. Ridvan Kiral, Suchetana B. Dutta, Gerit Arne Linneweber, Caroline Poppa, Max von Kleist, Bassem A. Hassan, Peter Robin Hiesinger

## Abstract

Variability of synapse numbers and partners despite identical genes reveals limits of genetic determinism. Non-genetic perturbation of brain wiring can therefore reveal to what extent synapse formation is precise and absolute, or promiscuous and relative. Here, we show the role of relative partner availability for synapse formation in the fly brain through perturbation of developmental temperature. Unexpectedly, slower development at lower temperatures substantially increases axo-dendritic branching, synapse numbers and non-canonical synaptic partnerships of various neurons, while maintaining robust ratios of canonical synapses. Using R7 photoreceptors as a model, we further show that scalability of synapse numbers and ratios is preserved when relative availability of synaptic partners is changed in a DIPγ mutant that ablates R7’s preferred synaptic partner. Behaviorally, movement activity scales inversely with synapse numbers, while movement precision and relative connectivity are congruently robust. Hence, the fly genome encodes scalable relative connectivity to develop functional, but not identical, brains.

**One-Sentence Summary:** Non-identical connectivity and behavior result from temperature-dependent synaptic partner availability in *Drosophila*.

## Introduction

The genome encodes developmental programs that produce remarkably precise synaptic connectivity of intricate neural circuits. Yet, individual neurons across animal species, taken out of the context of these developmental programs, readily form ‘incorrect’ synapses, including with themselves (*1-5*). Even during normal development some degree of synaptic promiscuity is prevalent, e.g. as a basis for subsequent pruning or fine-tuning (*6-10*). The notion of promiscuous synapse formation is not at odds with precise outcomes. Instead, it offers the opportunity to explain precision in the context of developmental plasticity and robustness to perturbation. The limiting case of ‘total promiscuity’, i.e. the ability of any neuron to form synapses with any other neuron, is highly unlikely given known molecular interactions that specify or bias connections (*11-14*). At the other end of the spectrum, precise molecular key-and-lock mechanisms for all synapses represent the antithesis to promiscuous synapse formation: if the key does not fit the lock, a synapse should not form. This is equally unlikely, given the known ability, and often developmental necessity, to form synapses with variable partners. The developmental program can facilitate correct partnerships between neurons through restrictions of the time, location and kinetics of their interactions, but to what degree remains largely unresolved (*6*).

In genetically identical organisms brain wiring is not only precise, but also flexible, robust to perturbation, and variable within well-controlled limits (*6, 15, 16*). Non-genetic perturbation can therefore reveal the limits of genetic determinism when combined with a quantitative description of the limits of precision, flexibility and variability. A non-genetic perturbation that affects all developmental processes is temperature (*17, 18*). Animals have adopted one of two evolutionary strategies to ensure functional outcomes: either to precisely control the developmental temperature (e.g. human body or bee hive), or to evolve a developmental process that is temperature-compensated, i.e. robust to a temperature range (e.g. fish and flies). *Drosophila melanogaster* develops functional brains at temperatures between ∼15°C and ∼29°C, but it is unclear how these different developmental temperatures affect the precision of synaptic connectivity. Temperature strictly determines molecular kinetics apparent as Brownian motion. However, the extent to which subcellular dynamics, synapse formation and precise neural circuit formation are compensated for these molecular kinetic changes is, to our knowledge, not known for any neuron inside a developing brain. Similarly, to what extent connectivity-based behavior is in fact robust to non-genetic perturbation of development remains largely unknown.

Increasing temperature increases the pace of development in ectotherms such as amphibians and arthropods (*19-21*). Many neuron-based processes are temperature-compensated at the level of development or function, e.g. the precision of circadian clocks (*22*) and other rhythmic circuits (*23, 24*). On the other hand, developmental temperature can change outcomes, for example sex-determination in reptiles (*25, 26*). Already before 1920, studies in *Drosophila* revealed temperature-dependencies of the development of fly legs (*27*), wings (*28*) and eye facet numbers (*29*). More than 100 years later, the *Drosophila* connectome is being finalized based on specimens that developed at 25°C (*30-33*). It is unknown how the connectome might differ after development at 18°C.

In brain function, destabilizing variability is counterbalanced by synaptic scaling, a homeostatic change of many synapses that preserves the relative synaptic weight of each synapse (*34-36*). Temperature-dependent developmental scaling has been observed for the relative timing of events from cellularization through hatching in diverse *Drosophila* species (*21*). Developmental scaling of relative connectivity has, to our knowledge, not been documented.

In this study we quantitatively investigated the influence of developmental temperature on brain development from subcellular filopodial dynamics to synapse numbers, synaptic partnerships, neuronal branch complexity and behavior. We show that none of these parameters are fully compensated for developmental temperature. Instead, scaling of relative synaptic connectivity balances functional robustness with flexibility and variability of brain wiring. Specifically, lower temperature leads to an increase in synapse numbers and partnerships based on increased availability of axo-dendritic branches and filopodia. For R7 photoreceptor neurons we show that this increased availability leads to synapses with the same non-canonical synaptic partners as ablation of R7’s preferred postsynaptic partner. Synapse formation based on relative synaptic availability leads to stable ratios of majority synapses while total synapse numbers scale inversely with developmental temperature. Consequently, multiparametric behavior measurements in a visual choice assay revealed that movement precision and relative connectivity are congruently robust, while movement activity scales inversely with synapse numbers. Hence, evolution has selected for a *Drosophila* genome that can develop functional, but non-identical, brains through scalable synapse formation based on relative availability of synaptic partners during development.

## Results

*Drosophila* pupal development, the time during which the adult brain is wired, is almost precisely twice as fast at 25°C compared to 18°C based on common lab protocols and our recent measurement of wild type flies (2.04 faster, 98 hours vs 201 hours) (*37*). In theory, every molecular and cellular developmental process could be sped up by a factor of 2, which would result in identical outcomes after development at both temperatures (Fig.1A). If individual molecular or cellular processes are less than two-fold faster, compensatory feedback could keep the outcome robust to temperature variations. Alternatively, development at 18°C and 25°C could in fact lead to different outcomes. To quantitatively test how developmental processes and outcomes scale with developmental temperature, we devised a set of assays to track down temperature-dependencies from subcellular dynamics to behavior (Fig. 1A).

**Figure 1:**
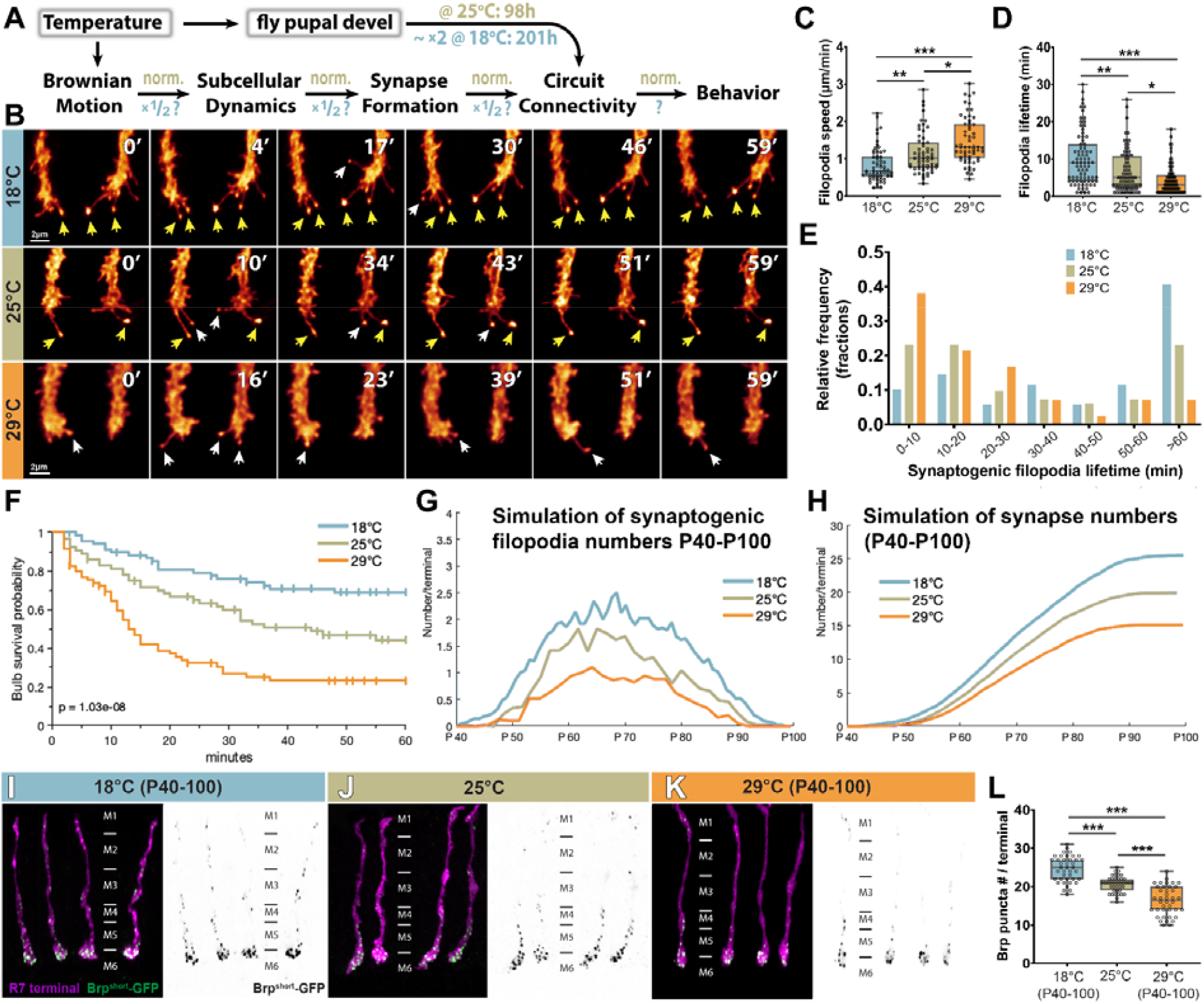
Temperature dependency of synapse formation in the R7 photoreceptor neuron. **(A)** Schematic of temperature effects on pupal developmental time and possible effects on subcellular dynamics, synapse formation, circuit connectivity and behavior. **(B)** Live imaging of filopodial dynamics during the time period of synapse formation (1 hour with 1 min time lapse; yellow arrows: long-lived synaptogenic filopodia; white arrows: short-lived synaptogenic filopodia). **(C)** Filopodial extension/retraction speeds are highly temperature-dependent. n=80 terminals per condition **(D)** Filopodial lifetimes are temperature-dependent. n=80 terminals per condition **(E)** The relative frequency of long-live synaptogenic filopdia (marked by bulbous tips) is temperature-dependent. n=23 terminals per condition. **(F-H)** Computational modeling predicts how the measured filopodia dynamics affect synapse formation. **(F)** Calculation of synaptogenic filopodia survival probabilities based on measure lifetimes that include filopodia that already existed at the beginning or still existed at the end of the imaging window. **(G)** Computational modeling of synapse development between P40-P100 based on synaptogenic filopodia dynamics. **(H)** Computational modelling of synapse number development between P40-P100 based on synaptogenic filopodia dynamics at different developmental temperatures. **(I-L)** Synapse numbers based on counts of the presynaptic active zone marker GFP-BrpD3 (Brp^short^) are dependent on the developmental temperature between P40-P100, the pupal time window when synapse formation occurs in the brain. n=40 terminals per condition. Data was analyzed with the Kruskal-Wallis test and Dunn’s as post-hoc test; *p<0.0332, **p<0.0021, ***p<0.0002.

### Subcellular dynamics and synapse formation of R7 photoreceptor neurons scale with developmental temperature

To measure the temperature-dependency of subcellular dynamics we first performed live imaging of developing R7 axon terminals during synapse formation, for which we have previously established quantitative dynamics (*38, 39*). First, we measured the temperature-dependency of filopodial dynamics that are known to quantitatively mediate synapse formation and partner choice (Fig. 1B) (*39-41*). At 18°C filopodia were 1.39-times slower and exhibited 1.52-times longer lifetimes compared to 25°C (Fig. 1C, D; Movie 1; Table S1).

These measurements reveal a temperature-dependency compared to a theoretical complete temperature compensation (factor of 1); however, these values indicate only partial temperature compensation relative to the increased speed of pupal development (factor of 2). Development at 29°C (at the upper end of tolerable fly developmental temperatures) further exacerbated this effect (Fig. 1C, D; Movie 1). The relative frequency of synaptogenic filopodia with longer lifetimes (marked by bulbous tips and increased stability) was similarly increased at lower temperatures (Fig. 1E). Hence, subcellular dynamics that underlie the development of synaptic connectivity differ significantly at different developmental temperatures.

Synaptogenic (bulbous) filopodia are the result of pre- and postsynaptic partner contacts based on our previous analysis of several mutants that selective affect contact initiation, stabilization and synapse maturation (*39*). Hence, the kinetics of synaptogenic filopodia throughout development allow to quantitatively predict adult synapse numbers, as previously shown in a computational model of this process (*39, 40*). To apply this model for different developmental temperatures, we first calculated the extrapolated real lifetimes of synaptogenic filopodia based on 60-minute live imaging data taking into consideration filopodia that were already present at the beginning or still present at the end of the imaging window. At 18°C, synaptogenic filopodia had a probability of 69% to live for at least 60 minutes, which reduced to 45% at 25°C and 23% at 29°C; synaptogenic filopodia lifetimes were 1.52-times higher at 18°C compared to 25°C (Fig. 1F; Table S1). Based on these filopodia lifetimes, a Markov state model simulation predicts the progression of synapse formation throughout pupal development; the model accurately recapitulates progression of synaptogenic filopodia occurances (Fig. 1G) and predicts significantly different synapse numbers after development at the three different temperatures (Fig. 1H; see Mathematical Modeling in Methods). To test these predictions, we measured synapse numbers immediately after pupal development at 18°C, 25°C or 29°C using three independent methods.

First, we used the presynaptic release site marker BrpD3 probe (*42*), which revealed significantly increased numbers following development at lower temperature (Fig. 1I-L).

Adult synapse numbers after development at the three different temperatures were in line with the model predictions (Fig. 1L; Table S1).

Second, to count synaptic connections based on synaptically connected postsynaptic partners of R7 neurons, we used a specific driver line that labels a subset of R7 neurons named yellow R7 (yR7) (reviewed in (*43*)) and the genetically encoded trans-synaptic tracer technique trans-Tango (*44*). Adult connectivity based on trans-Tango revealed a 1.26-times increase of postsynaptically connected partners after development at 18°C compared to 25°C (Fig. 2A-D; Table S1). Remarkably, the trans-Tango labeling reproducibly identified several cell types after development at 18°C that are very rarely, or not at all, postsynaptically connected to yR7 photoreceptors according to available connectome data (*31*). In particular, we found R7 synaptic connections to interneurons of the C2/C3 cell type (0 synapses in the connectome), Tm9 cells (0 synapses in the connectome), and Mi cell types including Mi1 and Mi4; 0-2 synapses in the connectome) (Fig. 2A; Fig. S1A). Notably, the electron microscopy-based connectome data is based on a specimen that developed at 25°C. Correspondingly, and consistent with the connectome data, these cells were not detected in trans-Tango experiments after development at 25°C or 29°C (Fig. 2B-D). These findings suggest increased synapse numbers after developmental at lower temperatures that may include synaptic partners excluded at higher developmental temperatures.

**Figure 2:**
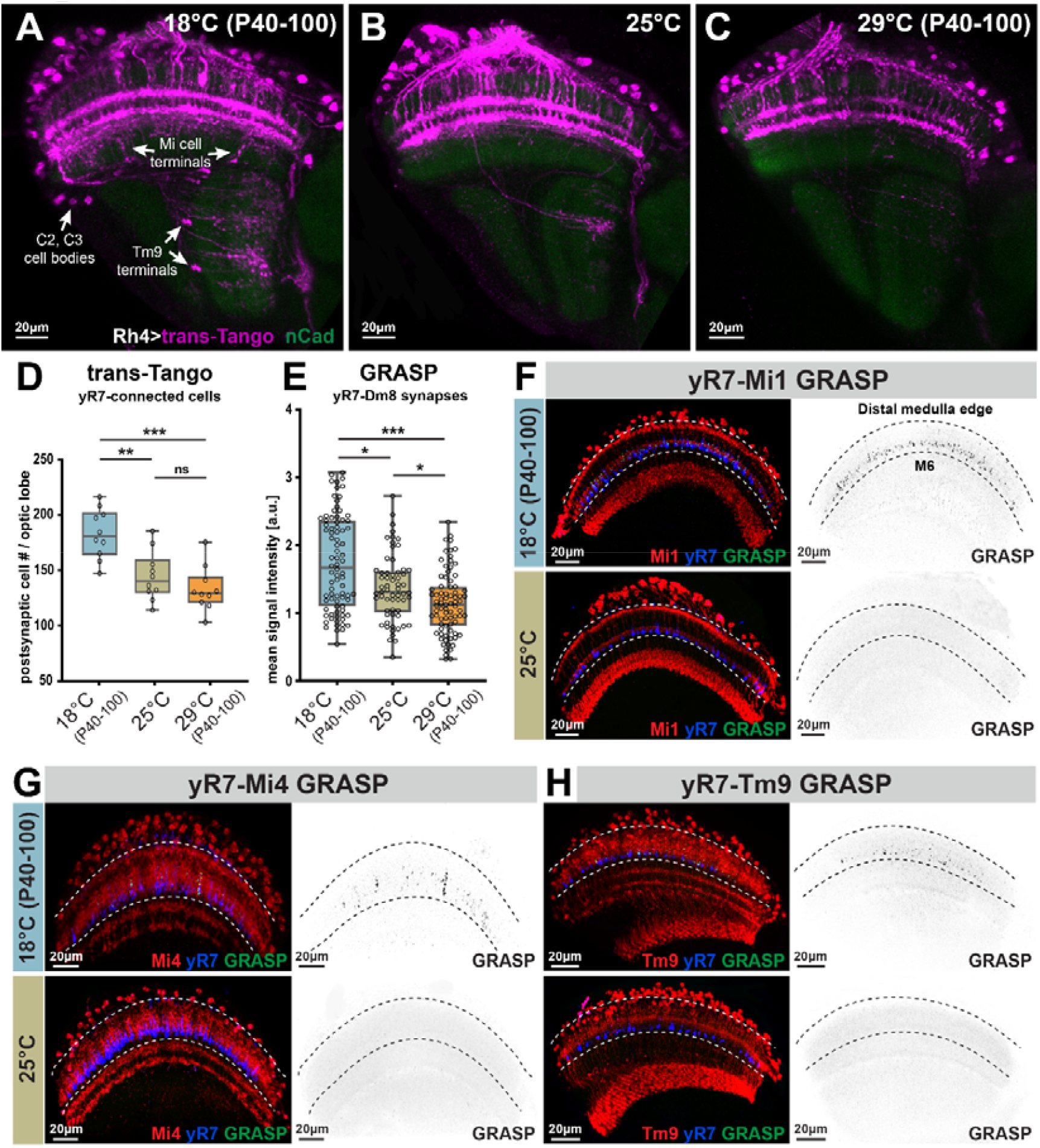
Synaptic connectivity scales with developmental temperature and includes additional, non-canonical synapses of yR7 photoreceptors at lower temperature. **(A-C)** Representative images of neurons (in magenta) that are postsynaptically connected to R7 neurons based on the genetically encoded trans-synaptic tracer method trans-Tango (*44*). See Figure S1A for identified cell types. **(D)** Quantification of cell body counts of trans-Tango labeled postsynaptic cells per optic lobe. n=10 optic lobes per condition. **(E)** Mean activity-dependent GRASP signal intensities between yellow R7 photoreceptors (yR7) and its main postsynaptic partner Dm8 neurons developed at different temperatures. n=85 terminals per condition. **(F-H)** Validation of active synapses using the activity-dependent GRASP method (*45*) for three identified postsynaptically connected neurons seen in **(A)** after development at 18°C that are not known to be synapically connected based on published connectome information (based on a specimen that developed at 25°C). Blue: yR7, Red: the potential postsynaptic partner, Green: activity-dependent GRASP signal (reconstituted GFP). Black/white panels: single channel of the green GRASP signal. Data was analyzed with the Kruskal-Wallis test and Dunn’s as post-hoc test; *p<0.0332, **p<0.0021, ***p<0.0002, ns=not significant.

Note that in all experiments only the developmental temperature during brain wiring between 40% and 100% of pupal development was varied; subsequently the newly hatched flies of all experimental groups were kept at 25°C for one week and treated identically for the trans-Tango labeling protocol. In contrast to developmental perturbation, alteration of the ‘functional’ temperature during the first week of life prior to the trans-Tango experiment did not lead to differences in the number of labeled postsynaptic cells (Fig. S2). Hence, our trans-synaptic tracing experiments suggest that synapse numbers are affected specifically by developmental temperatures and lead to differences of synaptic connections that remain stable for several days in the adult.

As a third method to validate changes of synapse numbers found in both BrpD3 active zone counts and trans-Tango experiments, we counted active synaptic connections using the activity-dependent GRASP method between specific synaptic pairs (*45*). We first test synapse numbers between yR7 and its main postsynaptic partner neuron, the amacrine-like cell type Dm8 (*46, 47*). Consistent with the connectome data, activity-dependent GRASP produced a strong signal precisely and selectively in the region where yR7 and Dm8 are known to forms synapses (Fig. S3). The synaptic labeling was 1.24-times stronger after development at 18°C compared to development at 25°C (Fig. 2E). This relative increase in synapse numbers based on activity-dependent GRASP is very similar to the increase in synapse numbers found using the presynaptic active zone marker (Fig. 1L) and trans-Tango (Fig. 2D). Finally, computational modeling based on measured filopodial dynamics yields the same numbers. We conclude that R7 synapse numbers inversely scale with developmental temperature based on four independent measurement methods.

Next, we tested whether the synaptic partnerships seen in trans-synaptic labeling after development at 18°C (but not at higher developmental temperatures) are in fact active synapses based on activity-dependent GRASP signals. Specifically, we tested interneurons Mi1, Mi4 and Tm9, for which the connectome analysis of a specimen that developed at 25°C have so far identified very few or no synapses. These are specifically: 2 R7-Mi1 synapses in one out of 18 reconstructed Mi1s; 1 R7-Mi4 synapse in one out of 16 reconstructed Mi4s; 0 R7-Tm9 synapses in 11 reconstructed Tm9s (*31*). We call these synapses ‘low-probability’ or ‘non-canonical’ synapses. Consistent with the connectome data, we found no GRASP signal between yR7 and Mi1, Mi4 or Tm9 after brain development at 25°C (Fig. 2F-H). By contrast, brain development at 18°C leads to robust labeling of these non-canonical synapses in exactly the layers where their axonal and dendritic processes are present (Fig. 2F-H). We conclude that development at 18°C leads to a significant increase of R7 synapses that includes both the main synaptic partner Dm8 as well as synaptic partners excluded during faster brain development at higher temperature.

We have previously shown that loss of autophagy in R7 photoreceptors leads to increased stability of synaptogenic filopodia and increased synapse formation with Dm8 as well as non-canonical postsynaptic partner neurons (*40*). The observation that lower temperature alone is sufficient to increase synapse formation with both canonical and non-canonical partner neurons suggests a model whereby all possible partner neurons increase their availability similarly, i.e. synaptic availability scales inversely with developmental temperature. For non-canonical partner neurons this model predicts a threshold effect, while maintaining relative synaptic ratios of canonical partner neurons (Fig. 3O). Synapse formation based on relative availability could in theory confer robustness to developmental temperature without keeping synapse numbers constant.

**Figure 3:**
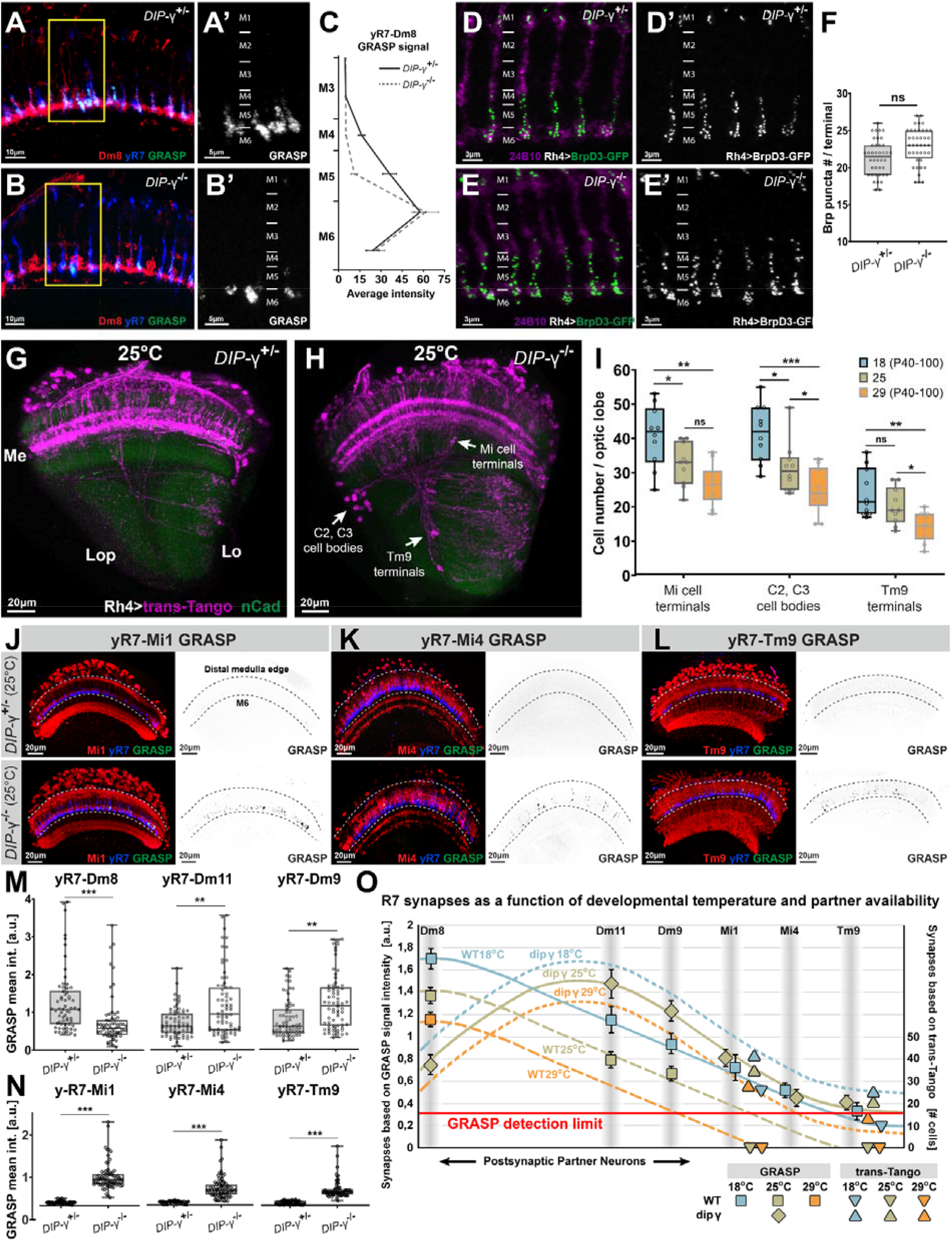
Loss of the postsynaptic partner neuron DIPγ(+)Dm8 reveals synapse formation of yR7 neurons based on relative availability. **(A-B)** Activity-dependent GRASP between yR7 and Dm8 neurons in a DIPγ mutant lacking DIPγ(+)Dm8 neurons and in a heterozygous control. **(A’-B’)** single channel GRASP signal. **(C)** GRASP signal intensity along the yR7 terminals reveals a loss of signal in the absence of DIPγ(+)Dm8 ‘sprigs’ between M4-M5, but no reduced intensity in the main synaptic layer M6. **(D-F)** Synapse numbers based on counts of the presynaptic active zone marker GFP-BrpD3 are not significantly altered in the DIPγ mutant. n=40 terminals per condition. **(G-H)** Representative images of the postsynaptically connected neurons (magenta) to yR7 based on the genetically encoded trans-synaptic tracer method trans-Tango. Note that the same additional cell types are present in the DIPγ mutant **(H)** as in wild type after development at 18°C (Fig. 2A). **(I)** The number of non-canonical postsynaptic partners in the DIPγ mutant is dependent on the developmental temperature; the mutant relative connectivity scales with temperature similar to wild type. n=10 optic lobes per condition. **(J-L)** Validation of active synapses using the activity-dependent GRASP method (*45*) for three non-canonical postsynaptically connected neurons identified in **(H)** in a DIPγ mutant lacking DIPγ(+)Dm8 neurons and in a heterozygous control that developed at 25°C. Blue: yR7, Red: the potential postsynaptic partner, Green: activity-dependent GRASP signal (reconstituted GFP). Black/white panels: single channel of the green GRASP signal. **(M)** Mean signal intensities of activity-dependent GRASP between yR7 and canonical partners (Dm8, Dm9, and Dm11) in a DIPγ mutant lacking DIPγ^+^Dm8 neurons and in a heterozygous control. n=70-80 terminals per condition. **(N)** Mean signal intensities of activity-dependent GRASP between yR7 and non-canonical partners (Mi1, Mi4, Tm9) in a DIPγ mutant lacking DIPγ(+)Dm8 neurons and in a heterozygous control. n=70-80 terminals per condition. **(O)** A model based on GRASP and trans-Tango counts that predicts a threshold effect for synapse formation between R7 and non-canonical partners (Mi1, Mi4, and Tm9), while maintaining relative synaptic ratios of canonical partner neurons (Dm8, Dm9, and Dm11). Data was analyzed with the Kruskal-Wallis test and Dunn’s as post-hoc test; *p<0.0332, **p<0.0021, ***p<0.0002, ns=not significant.

### Loss of yR7’s main postsynaptic partner neuron increases relative availability of the same non-canonical partners as development at lower temperature in wild type

To test relative partner availability and its role in temperature-dependent scaling of synapse numbers, we devised an experiment to change the availabilities of canonical and non-canonical postsynaptic partner neurons of yR7 photoreceptors. The yR7 subtype constitutes 65% of the total R7 population and specifically expresses the immunoglobulin cell adhesion molecule Dpr11, while the second R7 subtype is Dpr11 negative.

Correspondingly, the matching postsynaptic Dm8 partner neuron expresses the interacting partner molecule DIPγ, while the non-matching Dm8 neurons are DIPγ-negative. Curiously, loss of this molecular interaction leads to death of the majority of DIPγ(+)Dm8 cells (*48*), yet, surprisingly, the synapse numbers in presynaptic yR7 terminals remain unaltered in a *dpr11* mutant (*49*). If yR7s maintain their synapse numbers in the absence of their major synaptic partner, what other synaptic partner neurons do yR7 terminals recruit, and does their recruitment scale with different developmental temperatures similar to wild type?

To answer these questions, we first analyzed synapses between yR7 and possible partner neurons in a DIPγ mutant. We first validated the previously reported widespread loss of (∼2/3 of) Dm8 cells in this mutant (Fig. S4A-C). Loss of DIPγ(+)Dm8 cells is specific to columns containing the yR7 subtype and easily recognized by the loss of a Dm8 distal protrusion, also called ‘sprigs’, in such columns (*47, 50*); the sprigs mark an extended region of synaptic contacts between R7 and Dm8 cells (Fig. S4D-E). Correspondingly, we find less activity-dependent GRASP signal between yR7 neurons and Dm8 cells in the region of the missing sprigs due to missing DIPγ(+)Dm8 cells (Fig. 3A-C). Note that we used a Dm8 cell driver that expresses in both Dm8 subtypes and in the absence of two-thirds of all DIPγ(+)Dm8 cells active synapses with DIPγ(-)Dm8 cells and remaining DIPγ(+)Dm8 cells are detectable. Indeed, the most proximal region of yR7 terminals exhibits levels of activity-dependent GRASP between yR7 and Dm8 that are indistinguishable from control even in the absence of the matched DIPγ(+)Dm8 in the home column (Fig. 3B-C). Dm8 neurons are amacrine-like interneurons that extend axo-dendritic branches across more than 10 columns; consequently, a yR7 axon terminus with a missing matched DIPγ(+)Dm8 neuron is not prevented from forming synapses with processes from neighboring Dm8s despite their lack of the DIPγ, suggesting that DIPγ is not required for synapse formation between yR7-Dm8.

Due to the reduced number of synapses in the sprig region, the activity-dependent GRASP analysis between yR7 and Dm8s suggests an overall reduction of synapses between these two cell types (Fig. 3C). However, synapse counts based on the presynaptic marker BrpD3 indicate that yR7 synapse numbers in the DIPγ mutant were not significantly altered (Fig. 3D-F), in agreement with previous synapse counts in the *dpr11* mutant yR7s (*49*). To identify other postsynaptic partners, we performed trans-Tango experiments in the DIPγ mutant (at 25°C developmental temperature and using a +/DIPγ heterozygote as control; Fig. 3G, H). Remarkably, trans-Tango labeling in the DIPγ mutant optic lobe after development at 25°C looked very similar to a wild type optic lobe after development at 18°C, prominently including postsynaptically connected C2/3 cells, Tm9 cells and Mi cells (comp. Fig. 3H and Fig. 2A). Indeed, a cell-by-cell comparison revealed identical postsynaptically connected neurons for 18°C wild type and 25°C DIPγ (Fig. S1A-B).

Our trans-synaptic tracing results suggest that loss of DIPγ increases the relative availability of non-canonical partner neurons in a manner similar to lower developmental temperature in wild type, i.e. by increasing the pool of possible postsynaptic partners. While in wild type the low-probability synapses are effectively removed by a threshold effect at 25°C or above, their overall increase in the DIPγ mutant allows to test for their relative frequency at different temperatures. We found that the number of trans-synaptically labeled low-probability Mi1/4-yR7, C2/3-yR7 and Tm9-yR7 synaptic connections could be dialed down at 29°C and dialed up at 18°C, while maintaining their relative frequency (Fig. 3I, Fig. S4F-G). These findings indicate robustness of relative frequency to different developmental temperatures without keeping total synapse numbers constant, similar to our observation for canonical synapses in wild type (Fig. 3O).

To validate that the non-canonical connections represent functional synapses, we performed activity-dependent GRASP experiments for yR7-Mi1, yR7-Mi4 and yR7-Tm9 in control and DIPγ mutants after development at 25°C. As in wild type (Fig. 2F-H) and in the available connectome data, the +/DIPγ heterozygote control exhibited no or very rare GRASP signals for these non-canonical synapses. By contrast, the non-canonical synapses were prominent in the DIPγ mutant in the correct layer of their known axo-dendritic overlap and looked virtually indistinguishable from the wild type GRASP signal after development at 18°C (Fig. 3J-L, comp. Fig. 2F-H). Similarly, activity-dependent GRASP analyses of cell types that are known to be synaptically connected to yR7 (Dm9 and Dm11 cells) revealed similar increases of synaptic labeling in the DIPγ mutant (Fig. S5A-G; Table S2). These findings indicate that loss of DIPγ(+)Dm8 cells creates a situation in which yR7 neurons recruit more synaptic partners from a pool of both canonical (Fig. 3M) and non-canonical (Fig. 3N) partners. The recruitment of additional synaptic partners appears to be specific to those that have dendritic arborizations in the correct medulla layer, since Dm3 and Dm6 cells never form synapses with yR7 in control of the DIPγ mutant (Fig. S5H-K). Despite these shifts in synaptic partnerships, the final number of yR7 synapses is not significantly different from wild type at the same developmental temperature (Fig. 3F, 1L). These findings suggest a presynaptic mechanism for the determination of synapse numbers independent of the types of postsynaptic partners, consistent with the presynaptic serial synapse formation model for R7 (*39*) as well as previous observations for R1-R6 photoreceptors (*5*).

In sum, our analyses of yR7 synapse formation as a function of developmental temperature and partner availability revealed that relative synaptic frequencies are robust to different developmental temperatures (Fig. 3O). By contrast, overall synapse numbers increase with lower temperatures, including low-probability synapses not observed at higher temperatures. The robustness of relative synaptic frequencies to temperature as well as the total number of synaptic partners recruited by yR7 neurons is not affected by changes to the relative synaptic frequency itself. Specifically, loss of the majority of the main postsynaptic partner cells results in the same total number of synapses and the altered relative synaptic frequency scales with developmental temperatures similar to wild type. Notably, GRASP and trans-Tango measurements contribute in a consistent manner to a quantitative description of the temperature-dependency of relative connectivity (Fig. 3O); they are further consistent with both BrpD3 active zone counts and predictions of the computational model based on measured filopodial dynamics in wild type. Next, we set out to test whether these developmental temperature effects could be validated independent of genetic tools in functional and morphological measurements for a variety of neurons in the fly brain.

### Morphogenesis and synapse formation of branched interneurons in the brain depend on developmental temperature

To what extent is the temperature-dependency of neuronal development and synapse formation a general phenomenon in the fly brain? To approach this question, we analyzed a series of neuron types that face diverse challenges during the establishments of synaptic partner contacts.

First, we analyzed photoreceptors R1-R6, which terminate in a different brain region from R7, the lamina; in contrast to R7, growth cones of R1-R6 need to undergo a lateral sorting process to form a functional visual map required for motion vision according to the principle of neural superposition (*51-53*) and their functional output can be estimated based on electroretinogram recordings (ERGs) (*54*). Similar to our findings for R7, synapse numbers of adult R1-R6 increased by a factor of 1.15 based on the presynaptic BrpD3 marker after development at 18°C compared to development at 25°C (Fig. 4A, Fig. S6A-D).

**Figure 4:**
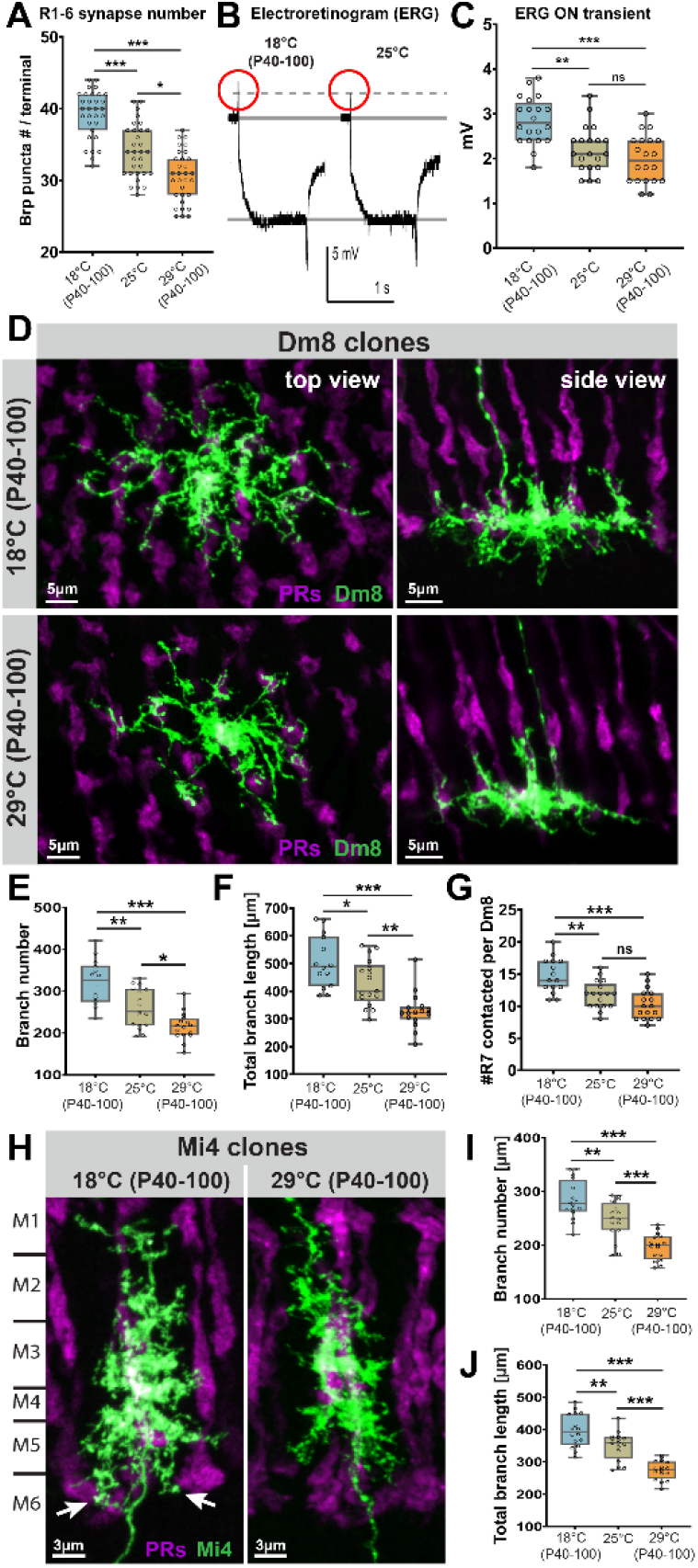
Neurotransmission of R1-R6 photoreceptors and branch morphology of Dm8 and Mi4 interneurons scale with developmental temperature. **(A)** R1-R6 photoreceptor synapse numbers based on counts of the presynaptic active zone marker GFP-BrpD3 (Brp^short^) are dependent on the developmental temperature between P40-P100, the pupal time window when synapse formation occurs in the brain. n=30 terminals per condition. **(B)** Representative electroretinogram (ERG) traces recorded from WT fly eyes developed at different temperatures. **(C)** Development at lower temperature increases neurotransmission of R1-R6 photoreceptors based on ERG ‘on-transient’ amplitudes. n=20 flies per condition. **(D)** Single cell clone representative images of Dm8 neurons developed at low (18°C, P40-100) and high (29°C, P40-100) temperatures. **(E-G)** Dm8 neurons exhibit increased branch numbers **(E)**, total branch length **(F)** and numbers of R7 contact sites **(G)** after development at 18°C. **(H)** Single cell clone representative images of Mi4 neurons developed at low (18°C, P40-100) and high (29°C, P40-100) temperatures. White arrows point Mi4 branches invading M6 medulla layer after development at 18°C where R7s are synaptically most active. **(I-J)** Mi4 interneurons (which are only connected to yR7 neurons after development at 18°C or in the absence of Dm8s) also exhibit increased branch numbers **(I)** and total branch lengths **(J)** after development at 18°C. Data was analyzed with the Kruskal-Wallis test and Dunn’s as post-hoc test; *p<0.0332, **p<0.0021, ***p<0.0002, ns=not significant.

Correspondingly, ERG recordings indicated that synaptic transmission (as measured by their ‘on’ transients) was significantly increased after development at 18°C compared to 25°C, consistent with increased numbers of synaptic connections (Fig. 4B-C, Fig. S6E, F). These findings indicate that the increased numbers of synapses after development at 18°C compared to 25°C are functional. By contrast, phototransduction, i.e. the ability of R1-R6 to convert a light stimulus into an electrical signal in the cell body (as measured by the ERG depolarization component) revealed no significant differences after development at different temperatures (Fig. S6E, G). We conclude that while phototransduction is fully compensated for variability of developmental temperatures, synapse numbers and synaptic output are increased at lower developmental temperatures consistent with our observations for R7 photoreceptors.

Next, we analyzed Dm8 neurons, the main synaptic partner neuron of R7 photoreceptors, whose dynamic and competitively regulated branch development has recently been analyzed in detail (*55*). Unexpectedly, the overall morphology of Dm8 neurons (Fig. 4D), including branch numbers (Fig. 4E) and total branch lengths (Fig. 4F) differed markedly depending on the developmental temperature. Since Dm8 extends its branches across several medulla columns, the temperature-dependency of its branch morphology leads to on average of 14 columns contacted by a single Dm8 after development at 18°C, 12 columns after development at 25°C and 10 columns after development at 29°C (Fig. 4G; Fig. S7A-C). These findings correspond well with the temperature-dependency of synapse numbers between yR7 and Dm8 (Fig. 2E; Table S1). The observations further suggest that increased branching of the canonical R7 partner Dm8 increases its availability for synapse formation.

In order to maintain robust relative synaptic frequencies, the synaptic availability of other neuron types needs to scale in a similar manner. To test whether increased availability through increased branching also occurs for a non-canonical partner of R7 photoreceptors, we analyzed Mi4 neurons after development at different temperatures. Similar to Dm8 neurons, Mi4 neurons exhibited a comparable temperature-dependency of both branch numbers and total branch lengths (Fig.4H-J; Table S1). Taken together, these findings suggest that interneuron branching, photoreceptor filopodial dynamics and synapse formation all scale with similar ratios with developmental temperature (Table S1).

### Dorsal Cluster Interneurons scale filopodial dynamics, branching, synapse numbers and synaptic partnerships with developmental temperature

To further analyze the generality and scalability of temperature-dependent branch extension and synapse formation, we focused on contralaterally projecting interneurons called dorsal cluster neurons (DCNs). DCNs form highly distinctive branched axonal patterns in the contralateral brain hemisphere (Fig. 5A); differences in these patterns predictively and quantitatively affect individual fly behavior (*15*). Similar to Dm8 and Mi4, we found that DCNs exhibited increased numbers of branches after development at 18°C compared to 25°C and 29°C (Fig. 5B-D). To observe the development of the different branching patterns, we established multiphoton live imaging of branching dynamics in the intact developing brain based on our *ex vivo* imaging culture system (*38*). Time lapse movies obtained during development at all three temperatures revealed a significant temperature-dependency of extension/retraction speeds, similar to R7 axon terminals (Fig. 5E-G; Movie 2; comp. Fig. 1B-D and Movie 1). Correspondingly, synapse numbers based on the BrpD3 presynaptic marker were increased after development at 18°C compared to 25°C and 29°C (Fig. 5H-J). These findings suggest that decreased branch dynamics and increased branch morphology lead to increased synapse numbers.

**Figure 5:**
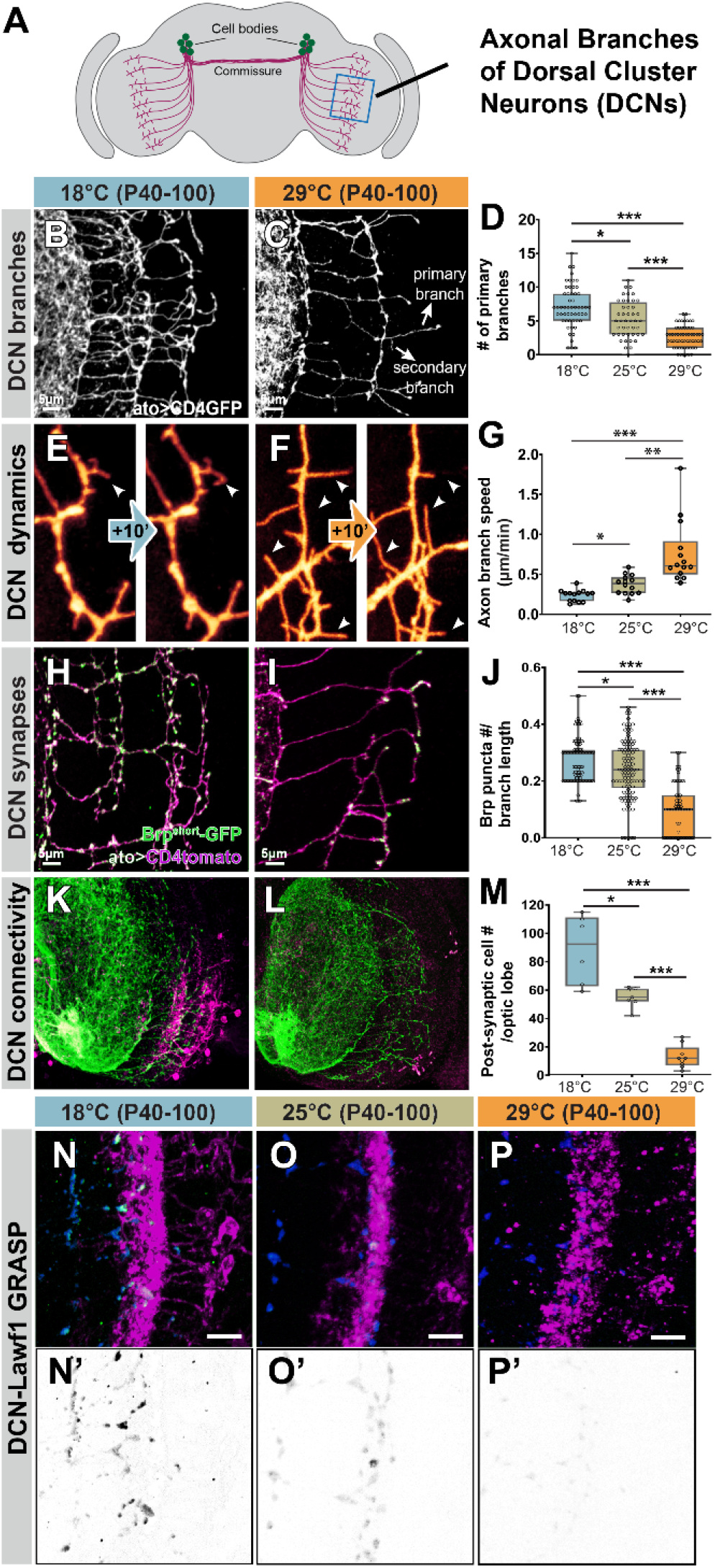
Branching dynamics, branch elaboration, synapse formation and partnerships of Dorsal Cluster Neurons scale with developmental temperature. **(A)** Dorsal Cluster Neurons (DCN) are large projection neurons with highly distinctive axonal branching patterns on the contralateral brain hemispheres. **(B-D)** DCNs exhibit increased number of axonal branches after development at 18°C. **(E-G)** Axonal branch dynamics of DCNs are highly temperature dependent. **(H-J)** Synapse numbers based on the presynaptic active zone marker GFP-BrpD3 reveals increased synapse formation of DCNs after development at 18°C. **(K-M)** DCNs form more postsynaptic connections after development at 18°C based on trans synaptic tracing method trans-Tango. **(N-P’)** Activity-dependent GRASP between DCN and Lawf2. Note increased GRASP signal after development at 18°C. Data was analyzed with the Kruskal-Wallis test and Dunn’s as post-hoc test; *p<0.0332, **p<0.0021, ***p<0.0002.

To validate the temperature-dependent scaling of synapse numbers and connectivity, we performed both trans-Tango and GRASP labelings of synaptic connections similar to our analyses of R7 neurons. Trans-synaptic tracing of postsynaptically connected cells with trans-Tango revealed a significant difference in the number of labeled postsynaptic cells after development at 18°C compared to 25°C, similar to our findings for R7 neurons (Fig. 5K-M). We identified several of the postsynaptically connected neurons based on morphology, including Lamina widefield (Lawf) interneurons as well as at least two more rarely connected cell types (L cells and Lpi cells) that we only observed after development at 18°C, but not 25°C (Fig. S7D-E). To validate the temperature-dependency of active synapses for these interneurons, we performed activity-dependent GRASP experiments between DCNs and Lawf1 neurons. We observed activity-dependent GRASP labeling of DCN-Lawf1 synapses after development at 18°C and to a significantly lesser degree after development at 25°C and 29°C; in all cases the GRASP signal was specific to the brain region where DCN-Lawf1 contacts are predicted (Fig. 5N-P). As with R7 photoreceptors, the temperature-dependent scaling of connectivity is supported by independent measurements of life dynamics and synapse numbers based on three independent methods. We conclude that slower development at lower temperature leads to decreased dynamics and increased branching and synapse formation with a larger pool of postsynaptic partners in DCNs. These findings further suggest that increased synaptic availability at lower developmental temperatures is a widespread phenomenon in the fly brain.

### Developmental temperature scales with movement activity, while movement precision is temperature-compensated

The developmental temperature-dependent differences of DCN branch morphologies are quantitatively comparable to differences in DCNs that are known to significantly affect behavior in a visual choice assay (*15*). This assay, Buridan’s paradigm, is a multiparametric single fly behavioral paradigm that allows to quantitatively measure more than 25 different parameters related to fly movements in response to defined visual stimuli (black bars on two sides of an arena, Fig. 6A) (*15, 56*). The assay is sufficiently sensitive to measure specific behavioral differences as a consequence of different DCN morphologies (*15*). We set out to test behavioral differences in flies that exhibit more widespread morphological and synaptic differences, including the DCNs, following development at different temperatures. We assessed 25 behavioral parameters that include measures related to overall activity (e.g. walking speed, pause lengths, overall distance per time, etc.), measures for movement angles or location that are independent of visual cues (e.g. the amount of turns taken or the time spent at or away from the center of the arena), and measures directly related to movement angles or location relative to the visual cues (Table S3). Following development between P40-P100 at different temperatures, flies from all experimental groups were kept for one week at 25°C and behaviorally tested at 25°C. We tested a Janelia wild type strain used for recent connectome analyses (*31*).

**Figure 6:**
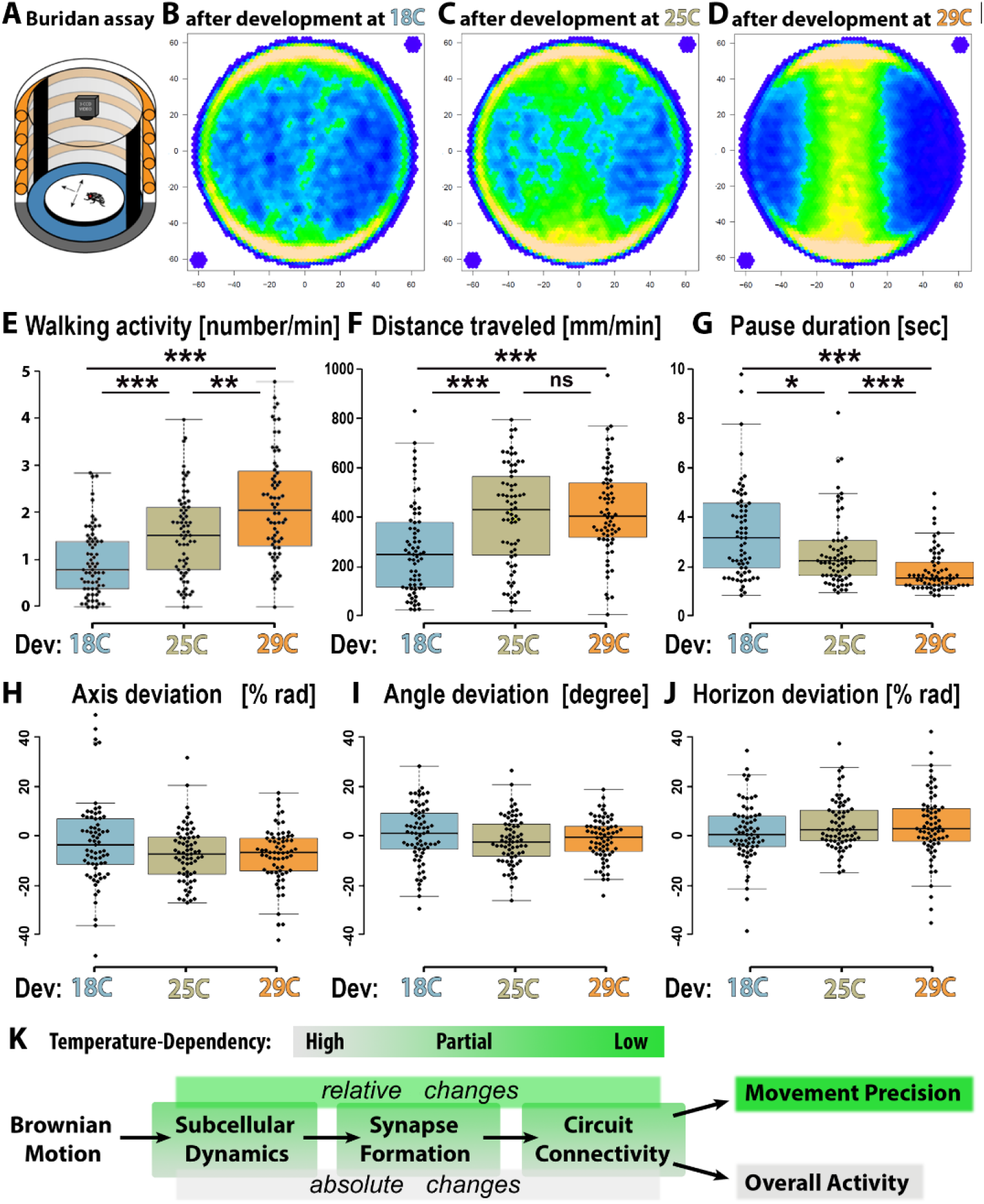
Movement activity scales with developmental temperature, while movement precision is temperature-compensated. **(A)**Schematic of Buridan’s paradigm. **(B-D)** Representative average traces of fly movements after development at different temperatures. **(E-J)** Six parameters of a total of 25 behavioral parameters for differences between adult behavior after development at 18°C, 25°C or 29°C. Quantitative data in Table S2. **(E-G)** The three most significantly temperature-dependent behavioral paramrters: Walking activity **(E)**, Distance Traveled **(F)** and Pause Duration **(G). (H-J)** Three fully temperature-compensated behaviors: axis deviation **(H)**, angle deviation **(I)** and horizon deviation **(J). (K)** Summary of temperature-dependencies. Subcellular dynamics, synapse formation and circuit connectivity are all temperature-dependent and partially temperature-compensated. By contrast, behavioral parameters associated with movement precision are fully temperature compensated, and behavioral parameters associated with overall movement activity are temperature-dependent. See Figure S8 for all behaviors and Materials and Methods for details on each parameter.

Development at 18°C, 25°C or 29°C led to significant behavioral differences that were immediately obvious from the paths and locations most frequented (Figs. 6B-D). Specifically, most parameters related to overall activity were significantly lower after development at 18°C compared to 25°C (Fig. 6E, Fig. S8), including number of walks (1,59 more; Fig. 6F), the total distance traveled (1,51 more; Fig. 6G) and pause durations (1,54 less; Fig. 6H). By contrast, none of the parameters associated with movement angles or location relative to the visual cues were significantly altered, e.g. axis deviation (Fig. 6I), angle deviation (Fig. 6J) or horizon deviation (Fig. 6K). Hence, parameters related to the precision of movement angles and relative connectivity were equally temperature-compensated, while general activity levels were highly temperature-dependent and scaled with developmental temperature in a range similar to neuronal branch morphologies and synapse numbers (Table S1, S3). We conclude that increased brain complexity (based on increased branch morphologies, synapse numbers and partnerships) is associated with reduced overall walking activity without affecting movement precision in Buridan’s paradigm in the temperature range between 18°C and 25°C. At the borderline physiological temperature of 29°C we further measured degradation of movement precision, presumably because of effects resulting from overall lower synapse numbers (Table S3).

In sum, we found significant behavioral differences after development at different temperatures, but also a fundamental ability of the *Drosophila* brain to compensate vitally important behavioral parameters for differences in brain wiring. Our findings suggest that robustness to variable developmental temperatures is achieved through scaling relative connectivity based on relative synaptic availability. These observations are consistent with evolutionary selection for functional flies, but not identical brains.

## Discussion

Neural circuits evolved based on genomes that contain information for the development, not the endpoint description, of functional connectivity. Selection occurs at the level of functional behavior as well as at the level of developmental robustness to variability of environmental conditions (*16*). Many animals, including *Drosophila*, have evolved robustness of brain development to varying developmental temperatures. However, robustness does not need to ensure identical development or outcomes at different temperatures, as long as the resulting connectivity is functional. In this study, we have shown that non-identical functional connectivity and adult behavior result from development at different temperatures. The underlying developmental processes do not specify synaptic connectivity in absolute terms, but based on scalable, relative availabilities of synaptic partners.

### Temperature robustness through developmental synaptic scaling

Developmental robustness could in theory result from compensation at a specific level, e.g. precision of synapse formation despite variable molecular and subcellular dynamics. In contrast, we found temperature-dependencies at every level from subcellular dynamics to synapse formation and circuit connectivity. However, the doubling of the developmental pace at 25°C compared to 18°C are not accompanied by a doubling of synapse formation. Instead, processes ranging from filopodial dynamics to branching and active zone formation were all only increased between ∼1.2-1.8-fold (Table S1). Consequently, development in half the time with less than a doubling of synapse formation leads to fewer overall synapses at 25°C compared to 18°C. In addition, both canonical and non-canonical synapses scale with developmental temperature, resulting in robust synaptic ratios, or relative connectivity, for majority synapses. In brain function, synaptic scaling is well characterized as a means of homeostatic regulation of destabilizing variability (*34*), with important consequences for learning and mental health (*35, 36*). The developmental synaptic scaling described here does not require the type of feedback mechanism underlying functional synaptic scaling (*34*); however, like its functional counterpart, developmental synaptic scaling provides as basis for the maintenance of relative input strengths in neural circuits.

We found that correct or incorrect relative connectivity scaled equally well with developmental temperature based on the R7 photoreceptor data. In the absence of their main postsynaptic partner, R7 neurons form more synapses with alternative partners to reach wild type synapse numbers. This altered synaptic connectivity scales in a temperature-dependent manner similar to wild type. It will be interesting to test these predictions for other neural circuits at the levels of individual neurons and circuit function, in particular in light of the recent development of a method to express any effector in precise proportions of neurons (*57*).

Every neuron type analyzed in this study exhibited a similar temperature-dependent scaling effect, including interneurons of the DCN, Dm8 and Mi4 type, as well as photoreceptors R1-6. Developmental parameters ranging from filopodial dynamics to branch numbers and length to synapse numbers scaled with temperature in a similar range and below the doubling rate required to compensate for a doubling of developmental pace at 25°C compared to 18°C (Table S1). These findings suggest that developmental synaptic scaling is a widespread phenomenon in the fly brain.

Similar to a lower temperature, reduced metabolism decreases the pace of *Drosophila* development and has recently been shown to increase robustness by decreasing developmental errors (*58*). While the mechanism of error suppression in this study is likely different from the temperature-effects observed here, we note that scalable relative connectivity is also likely more robust at lower temperatures because the synaptic ratio of high-probability synapses is established by larger numbers. In this context, it may be misleading to think of low-probability synapses as ‘errors’. Instead, temperature-dependent developmental scaling may add to concepts of wiring economy (*59-61*) by showcasing minimally sufficient connectivity after development at higher temperatures.

### A contribution of relative synaptic partner availability to synaptic connectivity

Lower developmental temperature in wild type flies increased numbers of synapses with both canonical and non-canonical partners, similar to previous observations in autophagy mutants (*40*). The increase in synapse numbers is mostly recruited from the most common synaptic partners, with rarer inclusion of postsynaptic partners that are excluded only at higher temperature. These findings suggest that temperature alone is sufficient to raise the availability of some synaptic partners above zero.

Recent connectome analyses based on specimens that developed at 25°C regularly find low-probability synapses that can plausibly be interpreted as errors (*30, 31*). On the other hand, our findings suggest that the number of these non-canonical synapses is significantly increased after development at lower temperature, without affecting the relative connectivity of high-probability synapses. Increased numbers of low-probability synapses correlate with reduced overall behavioral activity, but their functional significance remains unclear. Our live imaging and modeling of synapse formation suggest that increased stability of wild type filopodia and branches at lower temperature will robustly recruit non-canonical synaptic partners, similar to increased stability through a loss of developmental autophagy (*40*). Hence, temperature-dependent developmental synaptic scaling very likely requires proximity- and kinetics-based mechanisms involving locally restricted molecular machinery (*6, 62*). In addition, molecular specificity or selectivity with a ‘hierarchy of preference’ (*11*) are principally consistent with developmental synaptic scaling, as more stable filopodia and branches will also increase, and thus scale, molecular recognition. However, such a molecular ‘hierarchy of preference’ would have to include at least all the non-canonical synapses shown in this study for yR7 neurons and DCNs. To what extent molecular interactions play earlier developmental roles prior to synapse formation (*63*) and to what extent synapse formation could be promiscuous based on specification through proximity and kinetics (*6*) remains a matter of debate (*10, 11, 64*).

Our findings are consistent with the reported role of the Dpr11/DIPγ interaction for partner cell matching during a developmental process prior to synapse formation (*47, 48, 50*). The Dpr11/DIPγ interaction is required to ensure the survival and close proximity of a matched DIPγ(+)Dm8 cell to a Dpr11-positive yR7 terminal. Loss of DIPγ leads to loss of the majority of DIPγ(+)Dm8 cells and, as we show here, a subsequent widening of the pool of possible partners during the later process of synapse formation. The Drp11/DIPγ interaction effectively reduces the pool of postsynaptic partners by placing the main postsynaptic partner in close proximity and thereby increasing its relative availability.

Contrary to previous interpretations, we show that the subsequent synapse formation process does not utilize Dpr11/DIPγ interaction. Instead, yR7 axon terminals form a remarkably invariant number of synapses at a given temperature independent of the presence or absence of Dpr11, DIPγ, or the main postsynaptic partner neuron Dm8 (*49*)(this study). In a DIPγ mutant, yR7 axon terminals form synapses with non-matched Dm8s, plus other known synaptic partners based on the 25°C connectome, as well as available partners that are not, or very rarely, present in the 25°C connectome (*31*). Remarkably, the additionally recruited synaptic partners are identical in a DIPγ mutant after development at 25°C and wild type after development at 18°C. Hence, at least the cell types that are shown here to be recruited as postsynaptic partners during development at 18°C or in the absence of DIPγ(+)Dm8 cells, are not prevented from synapse formation by molecular mismatch. yR7 form an invariant number of synapses independent of what exact synaptic partners are more or less available.

Since both the number and the specificity of partnerships of yR7 synapses in the DIPγ mutant scale with developmental temperatures similar to wild type, we propose that synaptic specificity is a developmental outcome of a composite of relative contributors that include spatiotemporal availability, interaction kinetics, as well as interaction biases through molecular recognition between partner cells (*64*). In this view, the removal or alteration of a single relative contributor, e.g. the spatiotemporal availability of the main postsynaptic partner cell, increases the relative contribution of other factors, including the availability of other cells and their likelihood to form synapses based on interaction kinetics (*6, 40*).

### Temperature-dependent behavioral parameters - the philosopher and the chicken

The discovery of behavioral changes in response to environmental variability is reminiscent of the often excruciatingly stringent developmental conditions required for fly behavioral assays; typically, even small deviations from a specific developmental temperature preclude quantitatively meaningful behavior experiments with the adults. Based on our findings, these behavioral differences are a direct consequence of connectivity differences associated with different developmental temperatures.

Our observations suggest that temperature-dependent changes to brain wiring are a widespread phenomenon and not restricted to a specific neural circuit in the fly brain, because we found temperature-dependent changes for every developmental parameter and neuron type we analyzed. Our findings show that adult wild type flies that developed during the critical time period of synapse formation at a lower temperature have more elaborately branched interneurons, more synapses with more varied synaptic partners, and exhibit less overall movement activity. By contrast, adult flies that developed at the critical time period of synapse formation at a higher temperature have less branched interneurons, less synapses with fewer synaptic partners, and exhibit more overall movement activity. It is tempting to speculate about reduced brain connectivity underlying faster decision-making and more ‘frantic’ behavior, while increased brain complexity is commonly associated with more elaborate brain activity; however, the impact of these difference on behavioral fitness are unclear. Development at a certain temperature might prepare the adult for life at that temperature (*65*). Similarly, it is unclear whether ‘slower’ or ‘faster’ flies have a reproductive advantage when encountering one or the other type of fly. In either case, evolution is likely to have struck a balance between the selective advantages of developmental robustness to fluctuating temperatures and behavioral fitness.

## Acknowledgments

We would like to thank Emil Kind, Mathias Wernet, Claude Desplan and all members of the Hiesinger, Wernet and Hassan labs for their support and helpful discussions. We thank Mathias Wernet, Claude Desplan, Michael Reiser, Stephan Sigrist, Gilad Barnea, Ian Meinertzhagen, the Janelia Research Institute and the Bloomington *Drosophila* Stock Center for reagents. This work was supported by the NIH (RO1EY018884) and the German Research Foundation (DFG HI1886/5-1 and SFB186 TP02) and FU Berlin. B.H. was supported by an Einstein BIH Fellowship. M.v.K acknowledges financial support from the German ministry for education and science (BMBF) through grant number 01KI2016 and from the DFG, provided through the excellence cluster Math+, project EF3-2.

## Funding

National Institutes of Health grant RO1EY018884 (PRH)

German Research Foundation grant DFG HI1886/5-1 (PRH)

German Research Foundation grant SFB186 TP02 (PRH)

Einstein BIH Fellowship (BAH)

BMBF grant 01KI2016 (MvK)

Excellence Cluster Math+ (MvK)

## Author contributions

Each author’s contribution(s) to the paper should be listed [we encourage you to follow the <u>CRediT</u> model]. Each CRediT role should have its own line, and there should not be any punctuation in the initials.

Conceptualization: FRK, PRH

Methodology: FRK, SBD, CP, GAL, MvK

Investigation: FRK, SBD, CP, GAL, MvK

Visualization: FRK, SBD, GAL, BAH, MvK, PRH

Funding acquisition: BAH, MvK, PRH

Project administration: PRH

Supervision: BAH, PRH

Writing – original draft: FRK, PRH

Writing – review & editing: FRK, SBD, MvK, BAH, PRH

## Data and materials availability

All data are available in the main text or the supplementary materials.

## Supplementary Materials

Materials and Methods

Supplementary Text (Mathematical Modeling)

Figs. S1 to S8

Tables S1 to S3

References(##)

Movies S1 to S2

## Supplementary Materials for

## Materials and Methods

### Experimental model and subject details

Flies were reared at 25°C on standard cornmeal/yeast diet unless stated otherwise. For developmental analyses white pre-pupae (P+0%) were collected and staged to pupal developmental stages shown on figures. The following Drosophila strains were either obtained from Bloomington Drosophila Stock Center (BDSC) or other groups: UAS-Brp-short-GFP (S.Sigrist); Trans-tango flies (G.Barnea); DIPγ^null^ (C.Desplan); Rh4-Gal4, Rh4-LacZ (M.Wernet); R48A07-p65ADZp(attP40); R79H02-ZpGdbd(attP2) (Mi4-specific split Gal4 driver), Lawf1-Gal4 (M.Reiser); ato-Gal4-14a, ato-LexA, GRASP flies, hsflp, GMRflp, GMR-Gal4, GMR(FRT.stop)Gal4, FRT82B, GMR-Gal80, tub-Gal80, LexAop-CD8-GFP, UAS-CD4-tdGFP, UAS(FRT.stop)CD4-tdGFP, UAS-CD4-tdTomato, GMRmyrtomato, GMR49B06-LexA (Mi4-specific driver), GMR19F01-LexA (Mi1-specific driver), GMR25F10-LexA (Tm9-specific driver), GMR42H01-LexA (Dm9-specific driver), GMR20D11-LexA (Dm3-specific driver), GMR38H06-LexA (Dm6-specific driver), GMR11C05-LexA (Dm11-specific driver), ortC1-3-LexADBD, ortC2B-dVP16AD (Dm8-specific driver), GMR24F06-Gal4 (Dm8-specific driver) (BDSC).

### Drosophila genotypes

<u>Figure 1B-E</u>: GMRflp; GMR-Gal4, UAS-CD4tdGFP; FRT82B, tub-Gal80/FRT82B

<u>Figure 1I-L</u>: GMRflp; GMR-Gal4, UAS-CD4tdTomato/UAS-Brp^short^-GFP; FRT82B, tub-Gal80/FRT82B

<u>Figure 2A-D</u>: UAS-myrGFP, QUAS-mtdTomato(3xHA); Rh4-Gal4/*trans*-Tango

Figure 2E: Rh4-Gal4, UAS-nsyb::splitGFP1-10, LexAop-splitGFP11/ ortC1-3-LexADBD, ortC2B-dVP16AD (Dm8-specific driver)

<u>Figure 2F</u>: Rh4-Gal4, UAS-nsyb::splitGFP1-10, LexAop-splitGFP11/GMR19F01-LexA (Mi1-specific driver)

<u>Figure 2G</u>: Rh4-Gal4, UAS-nsyb::splitGFP1-10, LexAop-splitGFP11/GMR49B06-LexA (Mi4-specific driver)

<u>Figure 2H</u>: Rh4-Gal4, UAS-nsyb::splitGFP1-10, LexAop-splitGFP11/GMR25F10-LexA (Tm9-specific driver)

<u>Figure 3A-A’</u>: Rh4-Gal4, UAS-nsyb::splitGFP1-10, LexAop-splitGFP11/ortC1-3-LexADBD, ortC2B-dVP16AD (Dm8-specific driver); *DIPγ*^*null*^/+

<u>Figure 3B-B’</u>: Rh4-Gal4, UAS-nsyb::splitGFP1-10, LexAop-splitGFP11/ortC1-3-LexADBD, ortC2B-dVP16AD (Dm8-specific driver); *DIPγ*^*null*^/ *DIPγ*^*null*^

<u>Figure 3D-D’</u>: Rh4-Gal4/ UAS-Brp^short^-GFP; *DIPγ*^*null*^/+ <u>Figure 3E-E’</u>: Rh4-Gal4/ UAS-Brp^short^-GFP; *DIPγ*^*null*^/ *DIPγ*^*null*^

<u>Figure 3G</u>: UAS-myrGFP, QUAS-mtdTomato(3xHA); Rh4-Gal4/*trans*-Tango; *DIPγ*^*null*^/+

<u>Figure 3H</u>: UAS-myrGFP, QUAS-mtdTomato(3xHA); Rh4-Gal4/*trans*-Tango; *DIPγ*^*null*^/ *DIPγ*^*null*^

<u>Figure 3J:</u>Rh4-Gal4, UAS-nsyb::splitGFP1-10, LexAop-splitGFP11/ GMR19F01-LexA (Mi1-specific driver); *DIPγ*^*null*^/+ and *DIPγ*^*null*^/ *DIPγ*^*null*^

<u>Figure 3K:</u>Rh4-Gal4, UAS-nsyb::splitGFP1-10, LexAop-splitGFP11/GMR49B06-LexA (Mi4-specific driver); *DIPγ*^*null*^/+ and *DIPγ*^*null*^/ *DIPγ*^*null*^

<u>Figure 3L:</u>Rh4-Gal4, UAS-nsyb::splitGFP1-10, LexAop-splitGFP11/ GMR25F10-LexA (Tm9-specific driver); *DIPγ*^*null*^/+ and *DIPγ*^*null*^/ *DIPγ*^*null*^

<u>Figure 4A:</u>GMRflp; GMR-Gal4, UAS-CD4tdTomato/UAS-Brp^short^-GFP; FRT82B, tub-Gal80/FRT82B

Figure 4B-C: Canton-S WT flies

<u>Figure 4D-G</u>: hsflp; UAS(FRT.stop)CD4tdGFP; GMR24F06-Gal4 (Dm8-specific driver)

<u>Figure 4H-J</u>: hsflp; UAS(FRT.stop)CD4tdGFP/ R48A07-p65ADZp(attP40); R79H02-ZpGdbd(attP2) (Mi4-specific split Gal4 driver)

<u>Figure 5B-G</u>: ;UAS-CD4tdGFP/+;Ato-Gal4,UAS-CD4tdGFP/+ <u>Figure 5H-J</u>: ;UAS-Brp^short^-GFP/+;Ato-Gal4,UAS-CD4tdTomato/+

<u>Figure 5K-M</u>: UAS-myrGFP, QUAS-mtdTomato(3xHA); *trans*-Tango/+;Ato-Gal4/+

<u>Figure 5N-P’:</u> ;LexAop-nsyb::splitGFP1-10, UAS-splitGFP11/R52H01AD;Ato-LexA/R19C10DBD (Lawf1 split Gal4)

<u>Figure 6</u>: Canton-S WT flies (used in *(12)* to perform EM connectome analysis of synaptic partners in the *Drosophila* visual system)

<u>Supp. Fig. 1A</u>: UAS-myrGFP, QUAS-mtdTomato(3xHA); Rh4-Gal4/*trans*-Tango

Supp. Fig. 1B: UAS-myrGFP, QUAS-mtdTomato(3xHA); Rh4-Gal4/*trans*-Tango; *DIPγ*^*null*^/ *DIPγ*^*null*^

<u>Supp. Fig. 2:</u>UAS-myrGFP, QUAS-mtdTomato(3xHA); Rh4-Gal4/*trans*-Tango

<u>Supp. Fig. 3</u>: Rh4-Gal4, UAS-nsyb::splitGFP1-10, LexAop-splitGFP11/ ortC1-3-LexADBD, ortC2B-dVP16AD (Dm8-specific driver)

<u>Supp. Fig. 4A-C</u>: ortC1-3-LexADBD, ortC2B-dVP16AD (Dm8-specific driver), LexAop-CD8GFP

<u>Supp. Fig. 4D-D’</u>: ortC1-3-LexADBD, ortC2B-dVP16AD (Dm8-specific driver), LexAop-CD8GFP/Rh4-LacZ; *DIPγ*^*null*^/+

<u>Supp Fig. 4E-E’:</u>ortC1-3-LexADBD, ortC2B-dVP16AD (Dm8-specific driver), LexAop-CD8GFP/Rh4-LacZ; *DIPγ*^*null*^/ *DIPγ*^*null*^

<u>Supp. Fig. 4F-G</u>: UAS-myrGFP, QUAS-mtdTomato(3xHA); Rh4-Gal4/*trans*-Tango; *DIPγ*^*null*^/ *DIPγ*^*null*^

<u>Supp. Fig. 5B-B’</u>: Rh4-Gal4, UAS-nsyb::splitGFP1-10, LexAop-splitGFP11/GMR42H01-LexA (Dm9-specific driver); *DIPγ*^*null*^/+

<u>Supp. Fig. 5C-C’</u>: Rh4-Gal4, UAS-nsyb::splitGFP1-10, LexAop-splitGFP11/GMR42H01-LexA (Dm9-specific driver); *DIPγ*^*null*^/ *DIPγ*^*null*^

<u>Supp. Fig. 5E-E’</u>: Rh4-Gal4, UAS-nsyb::splitGFP1-10, LexAop-splitGFP11/GMR11C05-LexA (Dm11-specific driver); *DIPγ*^*null*^/+

<u>Supp. Fig. 5F-F’</u>: Rh4-Gal4, UAS-nsyb::splitGFP1-10, LexAop-splitGFP11/GMR11C05-LexA (Dm11-specific driver); *DIPγ*^*null*^/ *DIPγ*^*null*^

<u>Supp. Fig. 5H-H’</u>: Rh4-Gal4, UAS-nsyb::splitGFP1-10, LexAop-splitGFP11/GMR20D11-LexA (Dm3-specific driver); *DIPγ*^*null*^/+

<u>Supp. Fig. 5I-I’</u>: Rh4-Gal4, UAS-nsyb::splitGFP1-10, LexAop-splitGFP11/GMR20D11-LexA (Dm3-specific driver); *DIPγ*^*null*^/ *DIPγ*^*null*^

<u>Supp. Fig. 5J-J’</u>: Rh4-Gal4, UAS-nsyb::splitGFP1-10, LexAop-splitGFP11/ GMR38H06-LexA (Dm6-specific driver); *DIPγ*^*null*^/+

<u>Supp. Fig. 5K-K’</u>: Rh4-Gal4, UAS-nsyb::splitGFP1-10, LexAop-splitGFP11/ GMR38H06-LexA (Dm6-specific driver); *DIPγ*^*null*^/ *DIPγ*^*null*^

<u>Supp. Fig. 6A-D:</u>GMRflp; GMR-Gal4, UAS-CD4tdTomato/UAS-Brp^short^-GFP; FRT82B, tub-Gal80/FRT82B

<u>Supp. Fig. 6E-G</u>: Canton-S WT flies

<u>Supp. Fig. 7A-C</u>: hsflp; UAS(FRT.stop)CD4tdGFP; GMR24F06-Gal4 (Dm8-specific driver)

<u>Supp. Fig. 7D-E</u>: UAS-myrGFP, QUAS-mtdTomato(3xHA); *trans*-Tango/+;Ato-Gal4/+

<u>Supp. Fig. 8</u>: Canton-S WT flies

### Immunohistochemistry and fixed imaging

Pupal and adult eye-brain complexes were dissected in cold Schneider’s Drosophila medium and fixed in 4% paraformaldehyde (PFA) in PBS for 40 minutes. Tissues were washed in PBST (0.4% Triton-X) and mounted in Vectashield (Vector Laboratories, CA). Images were obtained with a Leica TCS SP8-X white laser confocal microscope with a 63X glycerol objective (NA=1.3). The primary antibodies used in this study with given dilutions were as follows: rat monoclonal anti-nCadherin (1:100; Developmental Studies Hybridoma Bank); goat polyclonal anti-GFP (1:1000; Abcam); rat monoclonal anti-GFP (1:500; BioLegend); rabbit polyclonal anti-CD4 (1:600; Atlas Antibodies); rabbit polyclonal anti-DsRed (1:500; ClonTech). The secondary antibodies Cy3, Cy5 (Jackson ImmunoResearch Laboratories) and Alexa488 (Invitrogen) were used in 1:500 dilution.

### Brain culture and live imaging

For all *ex vivo* live imaging experiments an imaging window cut open removing posterior head cuticle partially. The resultant eye-brain complexes were mounted in 0.4% dialyzed low-melting agarose in a modified culture medium. Live imaging was performed using a Leica SP8 MP microscope with a 40X IRAPO water objective (NA=1.1) with a Chameleon Ti:Sapphire laser and Optical Parametric Oscillator (Coherent). The excitation laser was set to 900 nm for single channel CD4-tdGFP imaging. Live imaging of R7 axon terminals at different temperatures was performed as follows: white pre-pupae (P+0%) were collected and staged to P+60% at 25°C. After eye-brain complexes were mounted in 0.4% dialyzed low-melting agarose in a modified culture medium, they were incubated 1 hour in imaging chamber at given temperatures on figures and scanned live for another hour with 1-min time resolution at the same incubation temperature. The same experimental flow and imaging settings were used for live imaging of Dorsal cluster neuron (DCN) axonal branches except that live imaging was performed at P+50%.

### Trans-tango and activity-dependent GRASP

Trans-tango and GRASP experiments were performed with yellow R7-specific driver Rh4-Gal4 and DCN-specific ato-Gal4-14a. Trans-tango flies were raised at 25°C until P+40% and moved to 18°C or 29°C for temperature shift experiments. On the day of eclosion, flies were transferred back to 25°C and dissected after 1 week. The number of postsynaptic neurons was counted manually from their cell bodies using cell counter plugin in Fiji including all cell bodies with weak or strong labelling to reveal all potential connections. Since postsynaptic partner labeling by Trans-tango is age-dependent, in another set of experiments, 1-day old flies were dissected to reveal the identity of cell types strongly connected to DCNs. For activity-dependent GRASP experiments, the same experimental flow was followed as in Trans-tango temperature shift experiments. To activate UV-sensitive R7-photoreceptors, flies were transferred to UV-transparent Plexiglas vials on the day of eclosion and kept in a custom-made light box with UV light (25°C, 20-4 light-dark cycle) for 4 days. To activate DCNs, flies were transferred to 25°C incubator with 12-12 light-dark cycle for 5 days. Brains were dissected and stained with a polyclonal anti-GFP antibody to label R7 photoreceptors, monoclonal anti-GFP antibody to label GRASP signal, and polyclonal anti-CD4 antibody to label postsynaptic neurons.

### Electroretinogram (ERG) recordings

Newly-hatched (0-day old) adult flies were collected and glued on slides using nontoxic school glue. Flies were exposed to alternating 1s “on” 2s “off” light stimulus provided by computer-controlled white LED system (MC1500; Schott). ERGs were recorded using Clampex (Axon Instruments) and quantified using Clampfit (Axon Instruments).

### Quantification and statistical analyses

#### Branch number and length analysis

All imaging data were analyzed and presented with Imaris 9.0.1 (Bitplane). Branches were detected automatically with the filament module using identical parameters for all experimental conditions (largest dendrite diameter: 3.0 µm, thinnest dendrite diameter: 0.2 µm). Inconsistencies in automatic detection were checked and corrected manually. The resultant values of branch numbers and lengths were taken and recorded directly from the statistics tab of the filament module. Graph generation and statistical analyses were done using GraphPad Prism 8.2.0

#### Synapse number analysis

All imaging data were analyzed and presented with Imaris 9.0.1 (Bitplane). For synapse number analysis, CD4-tomato channel was used to generate surfaces for individual R7 axon terminals and DCN axonal branches. Brp-positive puncta inside the surface were filtered using the masking function and were automatically detected with the spot detection module (spot diameter was set to 0.3 µ). Synapse numbers were taken and recorded directly from the statistics tab of the spot function. To obtain synapse density, the number of Brp-positive puncta inside individual DCN branch was divided by the respective branch length. Graph generation and statistical analyses were done using GraphPad Prism 8.2.0

#### Filopodia/axon branch tracing

Filopodia/axon branch tracing was performed using the filament module of Imaris 9.0.1 (BitPlane). Each filopodia/axon branch for all time points was segmented manually using “automatic placement” option to ensure the measurement of actual 3D length of each filopodia/axonal branch. Node was defined by the junction of axon shaft-branching point, from which filaments were created covering the entire length of the respective branch for all time points. “Length over time” data for all segmented filopodia/axon branch were recorded directly from the statistics tab of the filament module to calculate speed. Graph generation and statistical analyses were done using GraphPad Prism 8.2.0.

#### Statistical Analysis

Statistical comparison of two groups was performed with non-parametric Kolmogorov-Smirnov test. Statistical comparison of more than two groups was performed with non-parametric Kruskal-Wallis test and corrected for multiple comparisons with Dunn’s as a post-hoc test. All significance values are denoted on the graphs and in their respective legends. Graph generation and statistical analyses were done using GraphPad Prism 8.2.0.

#### Bulbous life time estimation

We used the Kaplan-Meier estimator provided in MatSurv (*66*) to estimate the Bulbous tip survival probability depicted in Fig. 1F.

### Buridan’s paradigm object orientation assay

Fly object orientation behavior was tested according to standard protocols in a Buridan arena (*15, 56*) using flies grown in a 12/12 h light–dark cycle. The arena consisted of a round platform of 117 mm in diameter, surrounded by a water-filled moat and placed inside a uniformly illuminated white cylinder. The assay was lid using four circular fluorescent tubes (Osram, L 40w, 640 C circular cool white) powered by an Osram Quicktronic QT-M 1 × 26–42. The fluorescent tubes were located outside of a diffuser (DeBanier, Belgium, 2090051, Kalk transparent, 180 g, white) positioned mm from the arena center. The temperature on the platform was 25 °C and 30 mm-wide stripes of black cardboard were placed on opposing sides inside of the diffuser. The retinal size of the stripes depended on the position of the fly on the platform and ranged from 8.4° to 19.6° in width (11.7° in the center of the platform). Fly tracks were analyzed using CeTrAn (*56*) and custom-written python code (*15*). We evaluated 44 partially overlapping behavioral parameters and have picked 25 representative from these for detailed analysis as shown in Fig. 6 and Supp. Fig. S8 and Table S2. The behavioral parameters are the following (*15, 56*):

## MEASURES OF OVERALL ACTIVITY

1. **Number of walks:** The number of times a fly walks from one stripe to the other. The fly needs to be on both ends near the edge more than 80% of the platform radius.
2. **Pause duration (s):** Median duration of pauses in seconds
3. **Distance traveled (mm/min):** Total distance travelled per minute.
4. **Relative time moving:** ratio of moving vs. not moving over the entire length of the fly track.
5. **Activity time (s):** Time active per minute in seconds
6. **Speed (mm/s):** Division of the distance traveled by time in mm/s. The reported value is the median speed of each fly. Movements exceeding 50mm/s are excluded in the median speed calculation.
7. **Number of pauses:** number of pauses per minute.
8. **Activity bouts (s):** Median duration of bouts of activity in seconds

## MEASURES OF MOVEMENTANGLES OR LOCATION INDEPENDENT OF VISUAL CUE

9. **Meandering (degrees/mm):** Measurement of the tortuosity (twistedness) of the track, calculated as Turning Angle divided by the speed. Shown as median value in degrees/mm.
10. **Turning angle (degrees)**: Median angle of all turns a fly does in the arena.
11. **Centrophobism while moving:** The arena is divided in an inner and outer ring of equal size. The ratio of time spend in the inner and outer ring is calculated. 1 signifies the fly has spent all its time in the outer part of the arena. -1 signifies the fly was at all times in the inner part of the arena. 0 would signifiy an equal distribution between inner and outer part of the arena: Only parts of the track while the fly is moving count to the calculation.
12. **Centrophobism while stationary:** Only parts of the track while the fly is not moving count to the calculation.
13. **Center deviation while moving:** Deviation away from the center of the platform. Values given in percent of the radius. Only parts of the track while the fly is moving count to the calculation.
14. **Center deviation while stationary:** Only parts of the track while the fly is not moving count to the calculation.

## MEASURES OF ANGLES OR LOCATION RELATIVE TO VISUAL CUE

15. **Absolute angle deviation:** Deviation angle from the path a fly walks away from the direction of the closest stripe. Direction does not matter. Median of all deviations is reported in degrees.
16. **Stripe deviation while moving:** Deviation away from the idealized line through the middle of the stripe. Direction towards right or left does matter. Values given in percent of the radius
17. **Stripe deviation while stationary:** Deviation away from the idealized line through the middle of the stripe. Direction towards right or left does matter. Values given in percent of the radius.
18. **Absolute stripe deviation while moving**: Deviation away from the idealized line through the middle of the stripe. Direction towards right or left does not matter. Values given in percent of the radius.
19. **Absolute stripe deviation while stationary:** Deviation away from the idealized line through the middle of the stripe. Direction towards right or left does not matter. Values given in percent of the radius.
20. **Angle deviation while stationary:** Deviation away from the idealized line through the middle of the stripe. Direction towards right or left does not matter. Values given in percent of the radius.
21. **Angle deviation while moving:** Deviation angle from the path a fly walks away from the direction of the closest stripe. Direction does matter. Median of all deviations is reported in degrees.
22. **Horizon deviation while moving:** Deviation away from the idealized line perpendicular to the stripes. Direction towards top or bottom stripe does matter. Values given in percent of the radius.
23. **Horizon deviation while stationary:** Deviation away from the idealized line perpendicular to the stripes. Direction towards top or bottom stripe does matter. Values given in percent of the radius.
24. **Absolute horizon deviation while moving:** Deviation away from the idealized line perpendicular to the stripes. Direction towards top or bottom stripe does not matter. Values given in percent of the radius.
25. **Absolute horizon deviation while stationary:** Deviation away from the idealized line perpendicular to the stripes. Direction towards top or bottom stripe does not matter. Values given in percent of the radius.

The data was statistically analyzed using the Kruskal-Wllis rank sum test and pairwise Wilcoxon rank sum test as a post-hoc test using R. (The post-hoc test was corrected with the Benjamini-Hochberg procedure to correct for multiple comparison.)

## Supplementary Text

### Mathematical Modeling

We adapted the data-driven stochastic model from Özel et al., 2019 by omitting the filopodia compartment and estimating temperature-specific parameters from the live imaging data (bulbous life time, number of bulbous tips at P60). In brief, we modelled synapses (S), short-lived transient bulbous tips (sB) and stable synaptogenic bulbous tips (synB).

The model’s reaction stoichiometries are determined by the following reaction scheme:

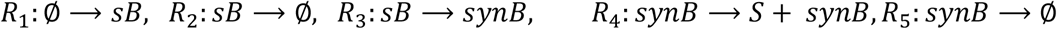

Reaction *R*_1_ denotes the formation of a (transient) bulbous tip, while *R*_2_ denotes its retraction. Reaction *R*_3_ denotes the stabilization of a transient bulbous tip, a stable bulb forms a synapse with reaction *R*_4_, while the bulbous tip remains visible and *R*_5_ denotes the retraction of a stable bulb. Note that in R_1_ we only implicitly model filopodia as outlined below.

Similar to the published model in Özel et al., 2019, reaction rates/propensities of the stochastic model are given by

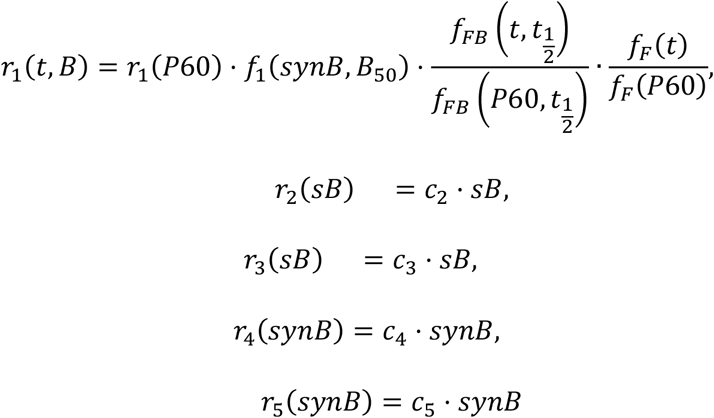

where *c*_2_,…, *c*_5_ are reaction constants (estimated as outlined below). The feedback function *f*_1_(*synB*, *B*_50_) = (*synB* + *B*_50_)/*B*_50_ models bulbous auto-inhibition due to limited resources and synaptic seeding factor competition as introduced before (Özel et al., 2019) and r_1_(*P*60) denotes the *net rate* of emergence of bulbous tips at *developmental time* P60. We do not consider the emergence of bulbous tips from filopodia as in previous work^1^, but rather implicitly through the time-dependent function *f*_*F*_(*t*). The functions *f*_*F*_(*t*) and *f*_*FB*_ 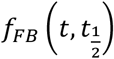) model slow-scale dynamics of filopodia- and bulbous dynamics, with previously determined parameters (Özel et al., 2019):

*f*_*FB*_(*t*) is a tanh function with

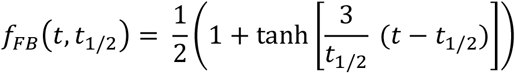

that models a time-dependent increase in the propensity to form bulbous tips with t_1/2_ = 1000 (min). The time-dependent function 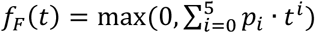 is a fifth-order polynome with coefficients p_5_ = -2.97 · 10^−14^, p_4_ = 3.31 · 10^−13^, p_3_ = -1.29 · 10^−9^, p_2_ = 2.06 · 10^−6^, p_1_ = -1.45 · 10^−3^ and p_0_ = 1 that down-regulates the generation of new filopodia at a slow time scale. Note, that *t* denotes the time in (min) after P40 (e.g. t_P40_ = 0 and t_P60_ = 60*20*scaling factor), which is scaled according to the factors discussed in Table MS1 below. The model was simulated using the Gillespie algorithm as outlined in (Özel et al., 2019).

### ‘Developmental time’ adjustment

At 25 °C, the pupal developmental stages correspond to the number of hours passed since pupation. For example, ‘P60’ refers to the pupal development stage observed at 60 hours past pupation at 25 °C. For the different temperatures the pupal development stages correspond to different durations past pupation. We measured that at 18 °C, the pupal development stage P100 is achieved 200.88 hours after pupation. At 29 °C it is achieved after 88.08 hours. Thus, for the different temperatures there are distinct factors that relate *real time* to *pupal development stage* as depicted in the Table below.

**Table MS1:** Factors that scale the ‘developmental stage’ to a ‘real time’ at different temperatures. We will use these scaling factors to relate *real time* to *developmental time* in our model simulations

| temp | developmental time P0 to P100 [hours] | factor |
| --- | --- | --- |
| 18°C | 200.88 | 2.05 |
| **25°C** | **98.16** | **1.00** |
| 29°C | 88.08 | 0.90 |

### Parameter estimation

Using the methods explained below, we derived the parameters depicted in Table MS2.

We first investigated the lifetimes of bulbous tips (Fig. 1F) and fitted parameter c_2_ and c_5_ which relates to the retraction of short- and long-lived bulbous tips, as shown in Figure MS1 below.

**Figure MS1:**
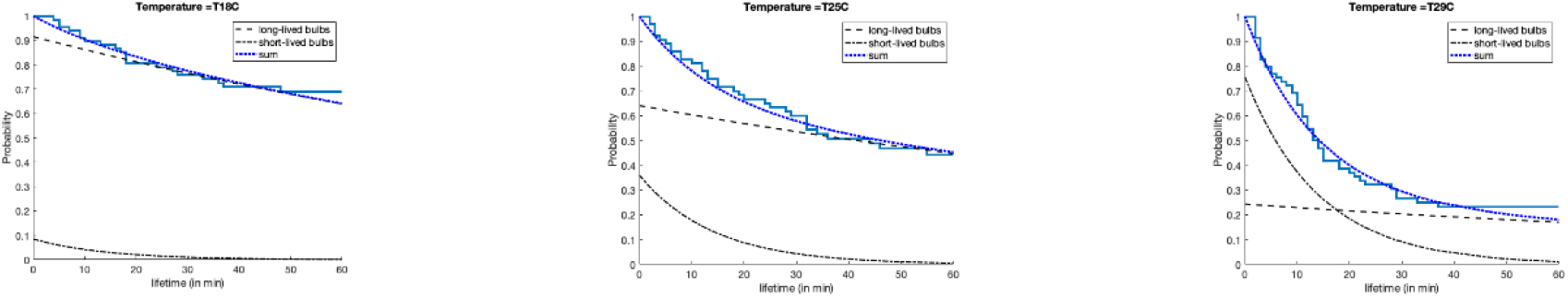
**Data derived (solid blue stairs) and fitted (continuous dotted blue line) bulbous lifetime kinetics. We assumed that the life time kinetics are resulting from the combined kinetics of short-lived and synaptogenic bulbous tips (black dotted and dashed lines). Fitted parameters: c_2_ = 0.0706 min^-1^; retraction rate of the synpatogenic bulbous tips was estimated to be c_5_ = 0.006 min^-1^ and the proportion of synaptogenic bulbous tips was 0.92, 0.64 and 0.24 at 18, 25 and 29°C.**

We then estimated the three parameters *r*_1_(*t* = *P*60), c_3_ and B_50_. To do so, we used the number distribution of short-lived and synaptogenic bulbous tips and set up the generator matrix

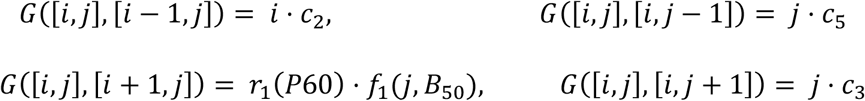

with diagonal elements such that the row sum equals 0. In the notation above, the tupel [i, j] denotes the state where *i* short-lived bulbous tips *sB* and *j* synaptogenic bulbous tips *synB* are present. The generator above has a reflecting boundary at sufficiently large N (maximum number of bulbous tips). Above, *r*_3_(*t*) is auto-inhibited by the number of stable bulbous tips through function *f*_1_. The stationary distribution of this model is derived by solving the eigenvalue problem

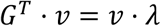

and finding the eigenvector corresponding to eigenvalue λ_0_ = 0. From this stationary distribution, we compute the marginal densities of *sB* and *synB* (e.g. summing over all states where i = 0, 1, … for sB) and fit them to the experimentally derived frequencies by minimizing the Kullback-Leibler divergence between the experimental and model-predicted distributions. Estimated parameters *r*_1_(*t* = *P*60), c_3_ and B_50_ are shown in Table MS2.

**Figure MS2:**
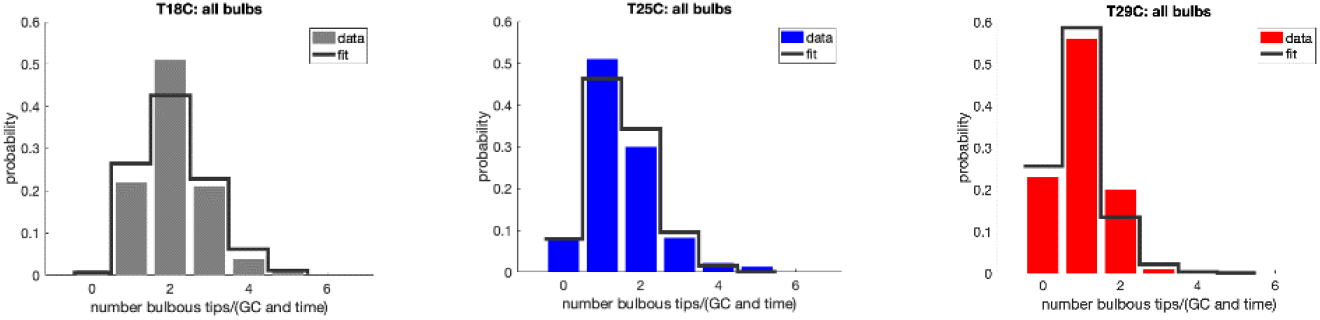
Data derived (bars) and fitted (solid line) number distribution for bulbous tips at developmental stage P60 for 18, 25 and 29°C. Finally, *c*_4_ was determined based on measured synapse numbers at P100.

**Table MS2:** Estimated parameters of the model. All parameters are in units min^-1^, except for B_50_ which is unitless.

| | $r_1(\text{P60})$ | $B_{50}$ | $c_2$ | $c_3$ | $c_4$ | $c_5$ |
| --- | --- | --- | --- | --- | --- | --- |
| 18°C | 0.3056 | 0.1187 | 0.0706 | 0.0427 | 0.0033 | 0.006 |
| 25°C | 0.0860 | 0.3505 | 0.0706 | 0.0207 | 0.0091 | 0.006 |
| 29°C | 0.0569 | 0.1044 | 0.0706 | 0.0081 | 0.0200 | 0.006 |

**Fig. S1.**
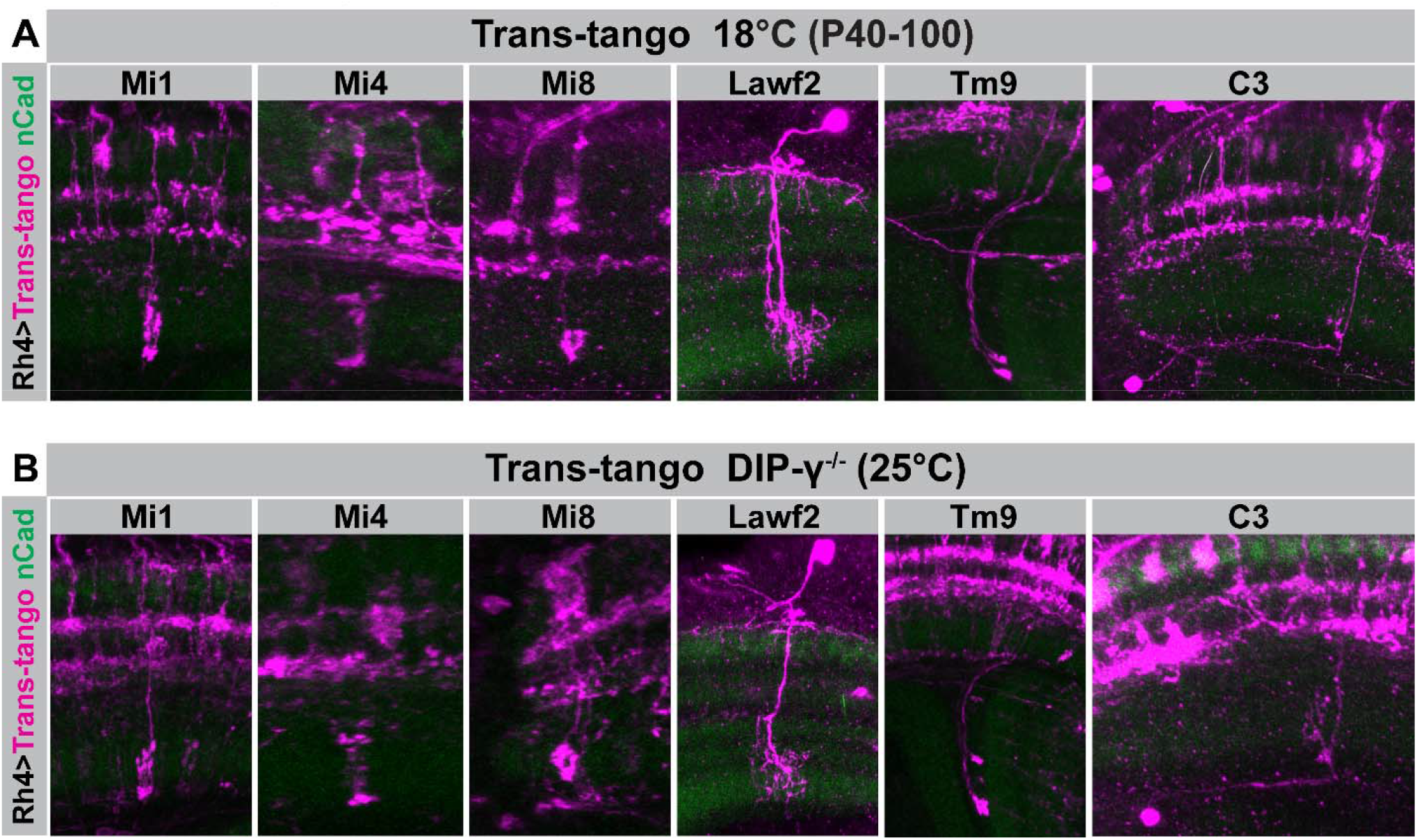
yR7s connect to the same types of non-canonical postsynaptic partners in wild type after development at 18°C and in DIPγ mutants after development at 25°C. **(A-B)** Representative images of neurons postsynaptically connected to yR7 photoreceptors in wild-type brains after development at 18°C **(A)** and in DIPγ mutant brains after development at 25°C **(B)**. Note that development at 18°C in wild type and at 25°C in the DIPγ mutant leads to non-canonical synaptic partnerships with the same type of neurons.

**Fig. S2.**
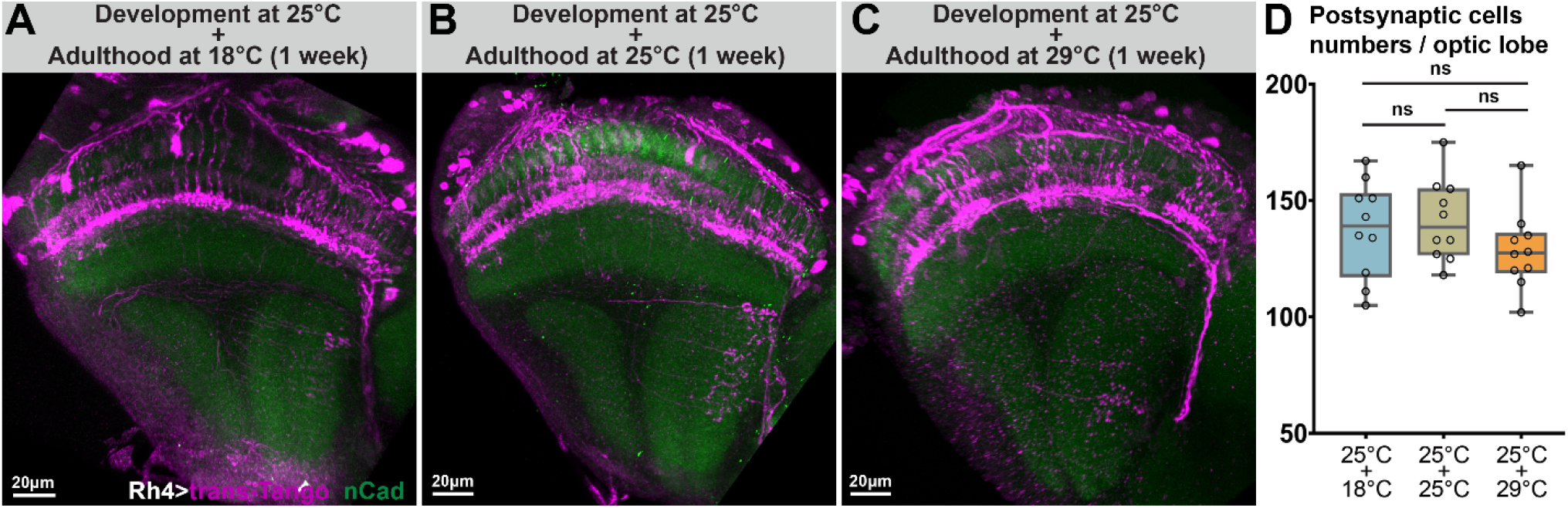
Alterations of temperature during the first week of life do not lead to differences in postsynaptically connected cell numbers. **(A-C)** Representative images of neurons postsynaptically connected to yR7 photoreceptors in brains developed at 25°C and then shift to either 18°C **(A)**, kept at 25°C **(B)** or shift to 29°C **(C)** during the first week of adult life prior to a trans-Tango experiments. **(D)** The number of neurons connected to yR7 photoreceptors do not significantly change when developed at the same temperature (25°C) and exposed to different ‘functional’ temperatures during adulthood. n=10 optic lobes per condition. Data was analyzed with the Kruskal-Wallis test and Dunn’s as post-hoc test; ns=not significant.

**Fig. S3.**
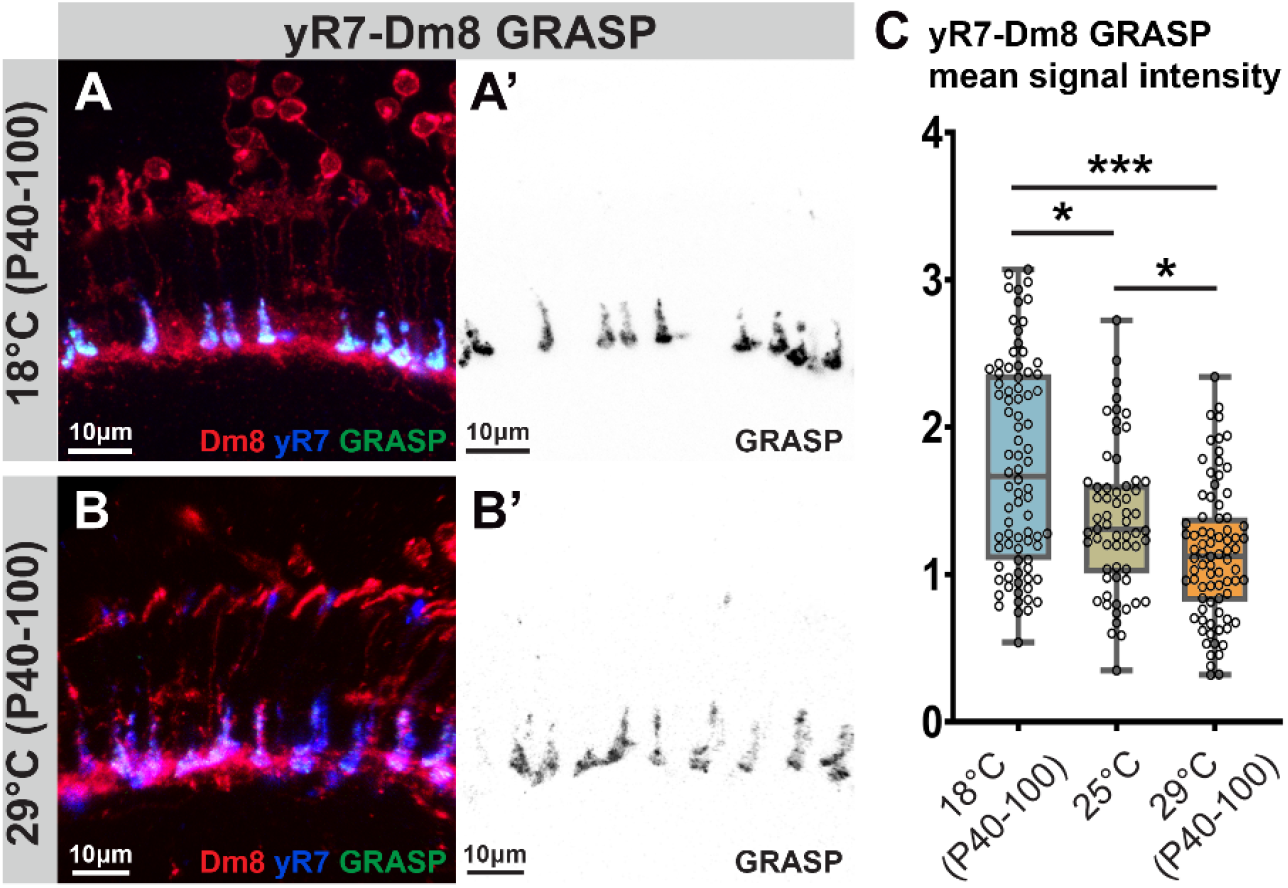
Activity-dependent GRASP labeling of yR7s and their main synaptic partner Dm8s scales with developmental temperature. **(A-B’)** Representative images of activity-dependent GRASP between yR7s and Dm8 after development at 18°C **(A-A’)** and after development at 29°C **(B-B’). (C)** Low developmental temperature scales with the GRASP signal between yR7s and their main synaptic partner Dm8. n=80 terminals per condition. Data was analyzed with the Kruskal-Wallis test and Dunn’s as post-hoc test; *p<0.0332, ***p<0.0002.

**Fig. S4.**
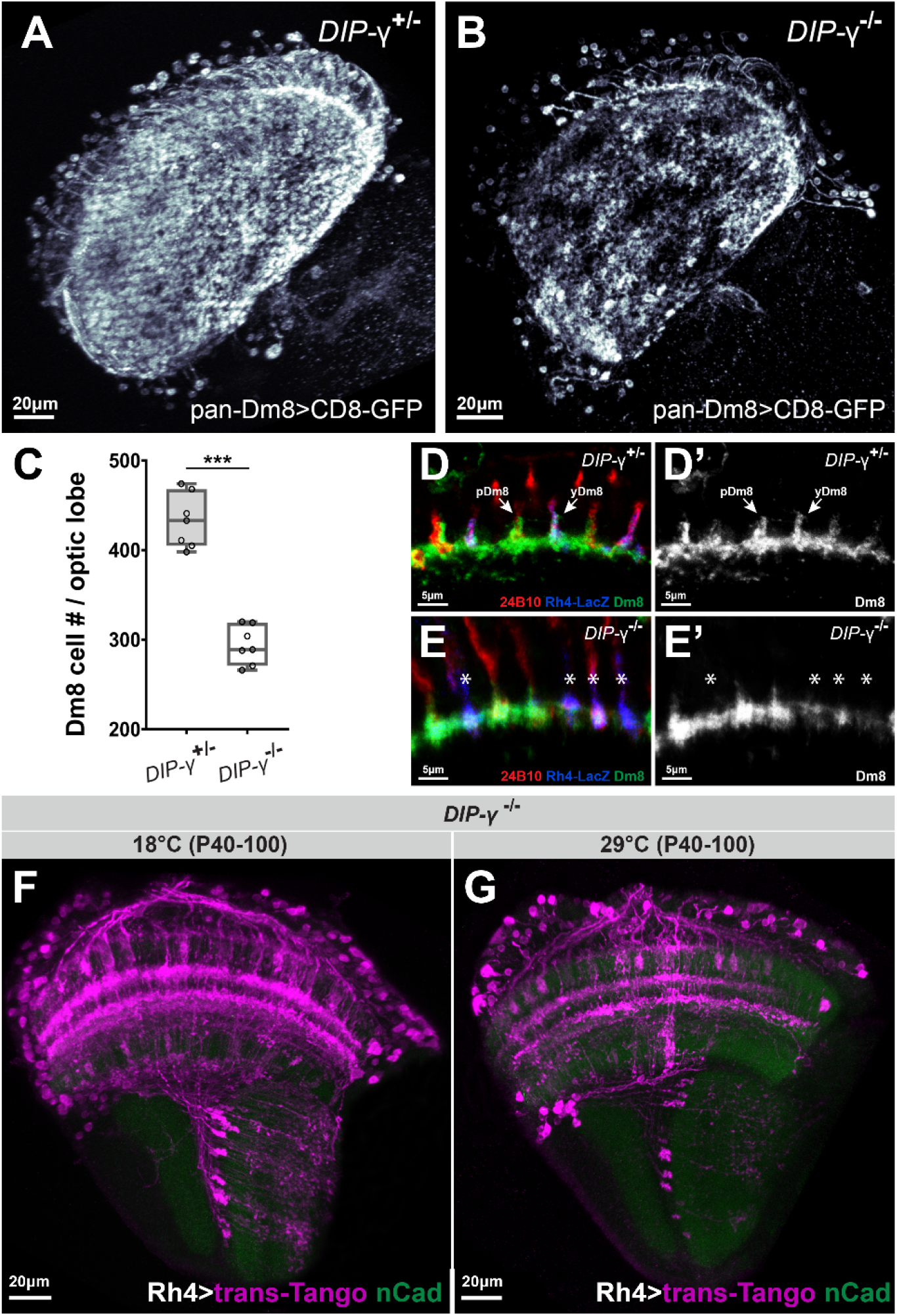
Loss of the majority of DIPγ^+^Dm8 neurons leads to increased recruitment of non-canonical synapses of yR7 neurons. **(A-C)** DIPγ loss-of-function leads to cell death of the majority of DIPγ^+^Dm8 neurons. n=7 optic lobes per condition. **(D-E)** Surviving DIPγ^+^Dm8 neurons lack distal membrane protrusions at medulla layers M4-M5. **(F-G)** Non-canonical partner neurons (Mi cells, C2/C3 cells, and Tm9) in DIPγ mutant brains scale with developmental temperatures of 18°C **(F)** and 29°C **(G)**, revealing an additive effect of low developmental temperature andmain synaptic partner loss on partner availability. Data was analyzed with the Kolmogorov-Smirnov test; ***p<0.0002.

**Fig. S5.**
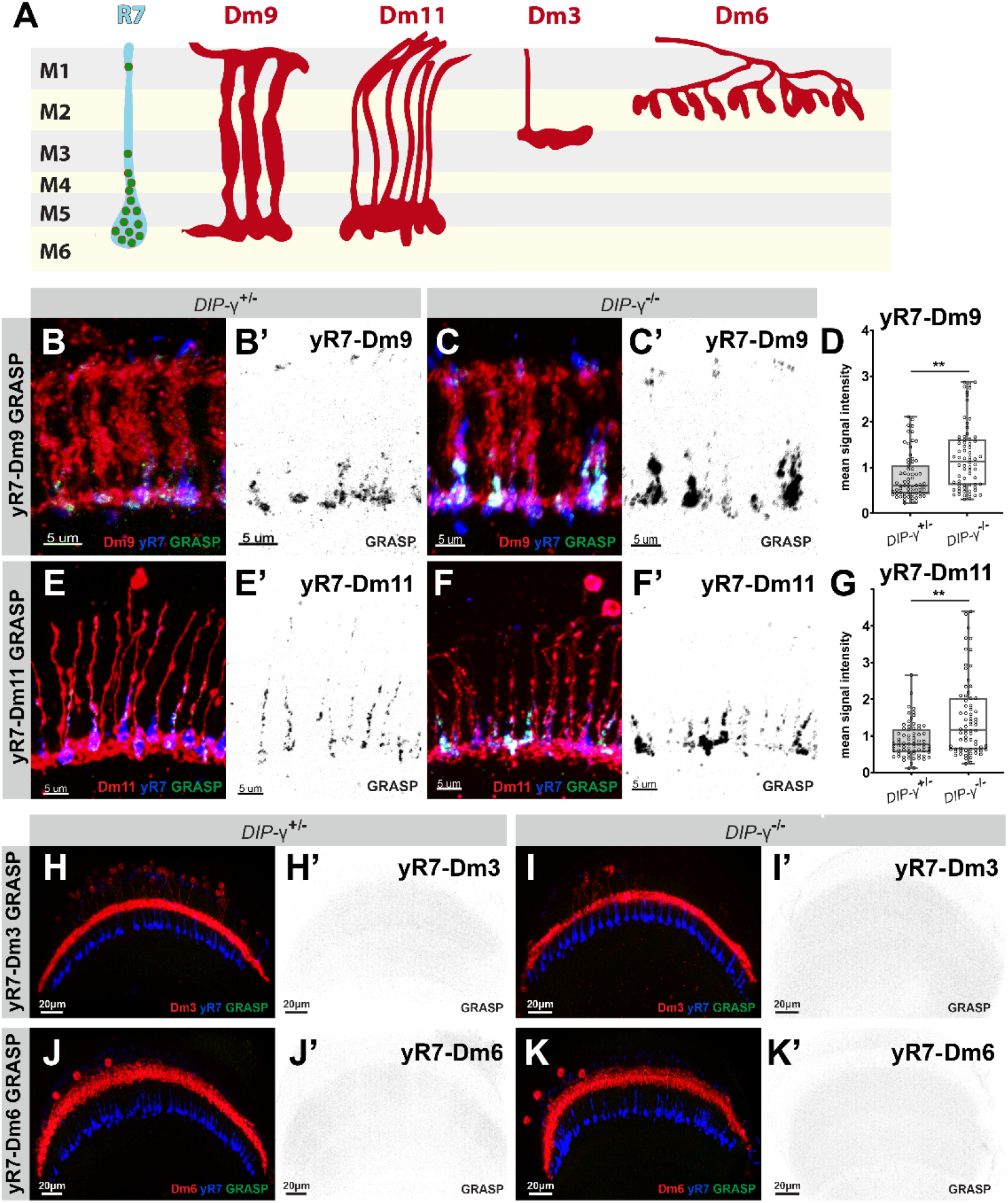
Loss of the majority of DIPγ^+^Dm8 neurons leads to increased activity-dependent GRASP signals of yR7 with its canonical synaptic partners Dm9 and Dm11. **(A)** Schematic representations of Dm9, Dm11, Dm6, and Dm3 neurons with an R7 terminal through medulla layers M1-M6. Green dots demonstrate the distribution of active zones in R7 terminals based on the connectome data. **(B-G)** Activity-dependent GRASP between yR7s and Dm9 **(B-D)** and between yR7s and Dm11 **(E-G)** in control and DIPγ mutant brains reveals stronger synaptic connections with the loss of main synaptic partner Dm8. n=60-80 terminals per condition. **(H-K’)** Dm8 neuron loss in DIPγ mutants does not lead to synapse formation with neurons (Dm3 and Dm6) that do not have dendritic branches in medulla layers. Data was analyzed with the Kolmogorov-Smirnov test; **p<0.0021.

**Fig. S6.**
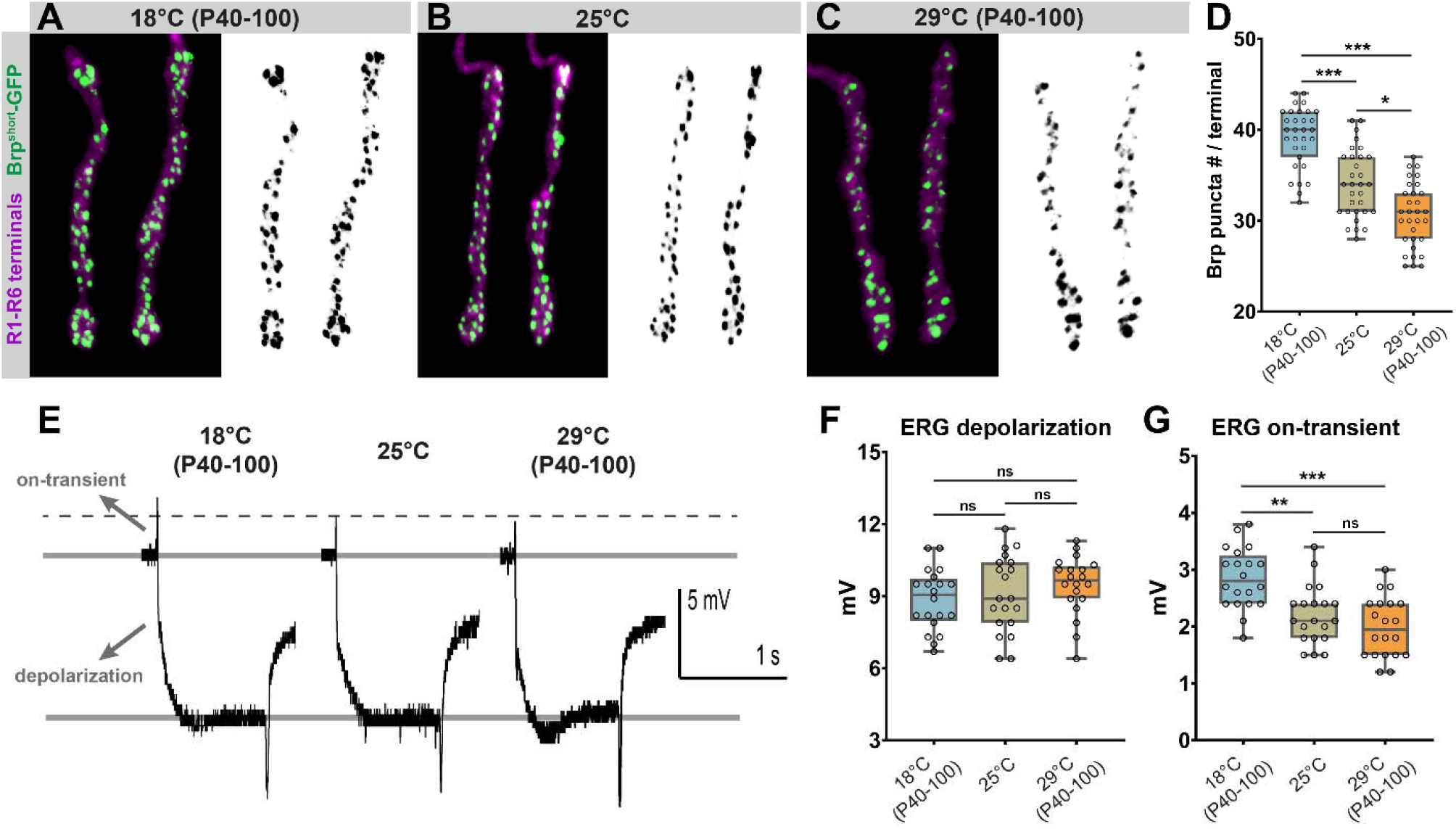
R1-R6 photoreceptor synapse number and neurotransmission scale with developmental temperature. **(A-C)** Representative images of R1-R6 photoreceptor axon terminals with GFP-BrpD3 (Brp^short^) marked active zones developed at 18°C **(A)**, 25°C **(B)**, and 29°C **(C). (D)** The number of active zones per R1-R6 terminal at different developmental temperatures. n=30 terminals per condition. **(E)** Representative electroretinogram (ERG) traces recorded from fly eyes developed at 18°C, 25°C, and 29°C. **(F)** Developmental temperature does not affect phototransduction based on ERG ‘depolarization’ amplitudes. n=20 flies per condition. **(G)** Low developmental temperature increases neurotransmission of R1-R6 photoreceptors based on ERG ‘on-transient’ amplitudes. n=20 flies per condition. Data was analyzed with the Kruskal-Wallis test and Dunn’s as post-hoc test; *p<0.0332, **p<0.0021, ***p<0.0002, ns=not significant.

**Fig. S7.**
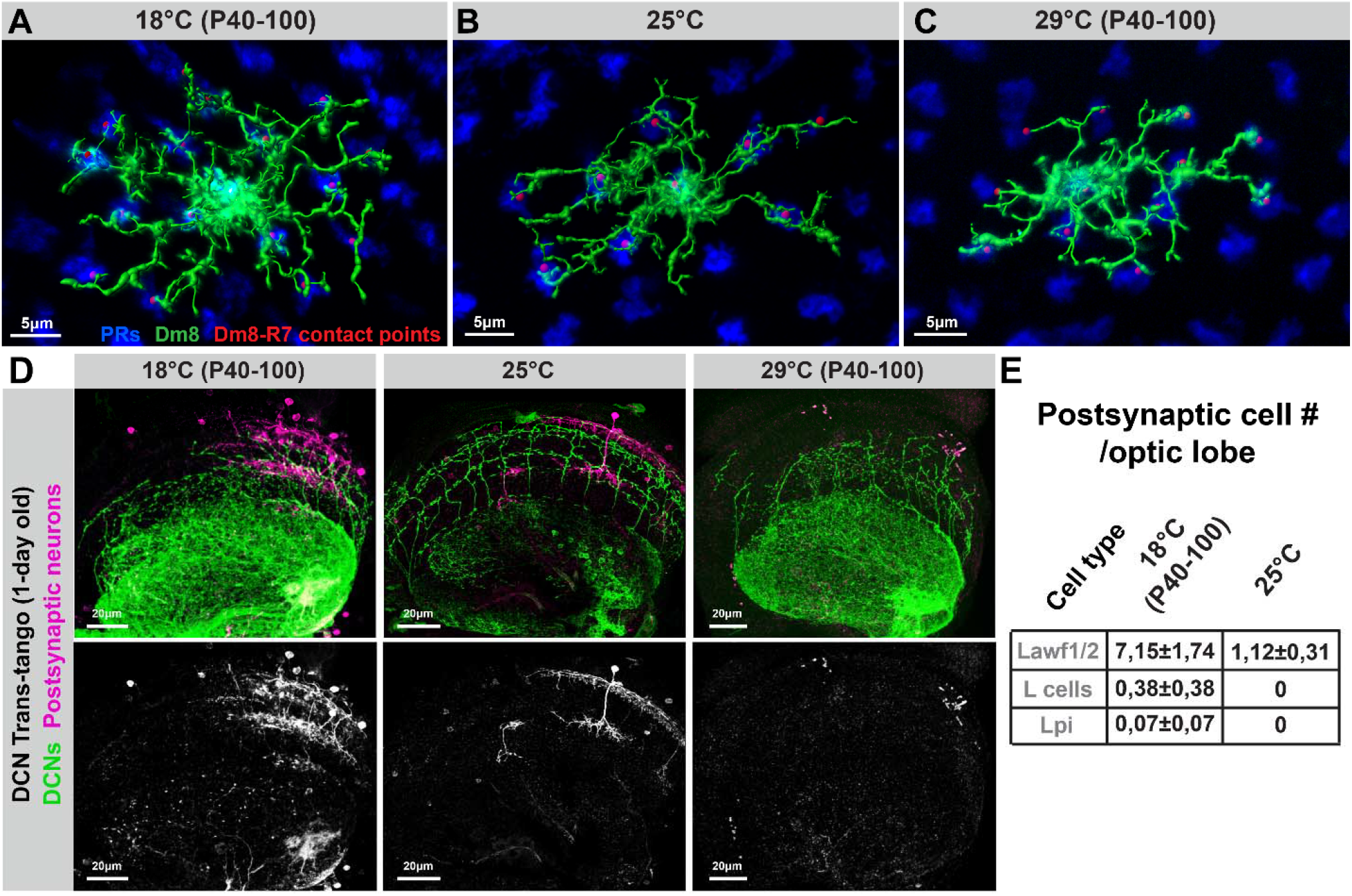
Branch elaboration and partner availability of Dm8s and DCNs scale with developmental temperature. **(A-C)** Skeleton reconstructions of Dm8 cells developed at 18°C **(A)**, 25°C **(B)**, and 29°C **(C)**. Red dots represent contact points between R7s and Dm8s. See Fig. 4E-G for corresponding quantifications. **(D-E)** Low developmental temperature leads to more widespread labeling of neurons postsynaptically connected to DCNs. Note that rarely connected cell types (L cells and Lpi cells) were only observed when brains developed at 18°C.

**Fig. S8.**
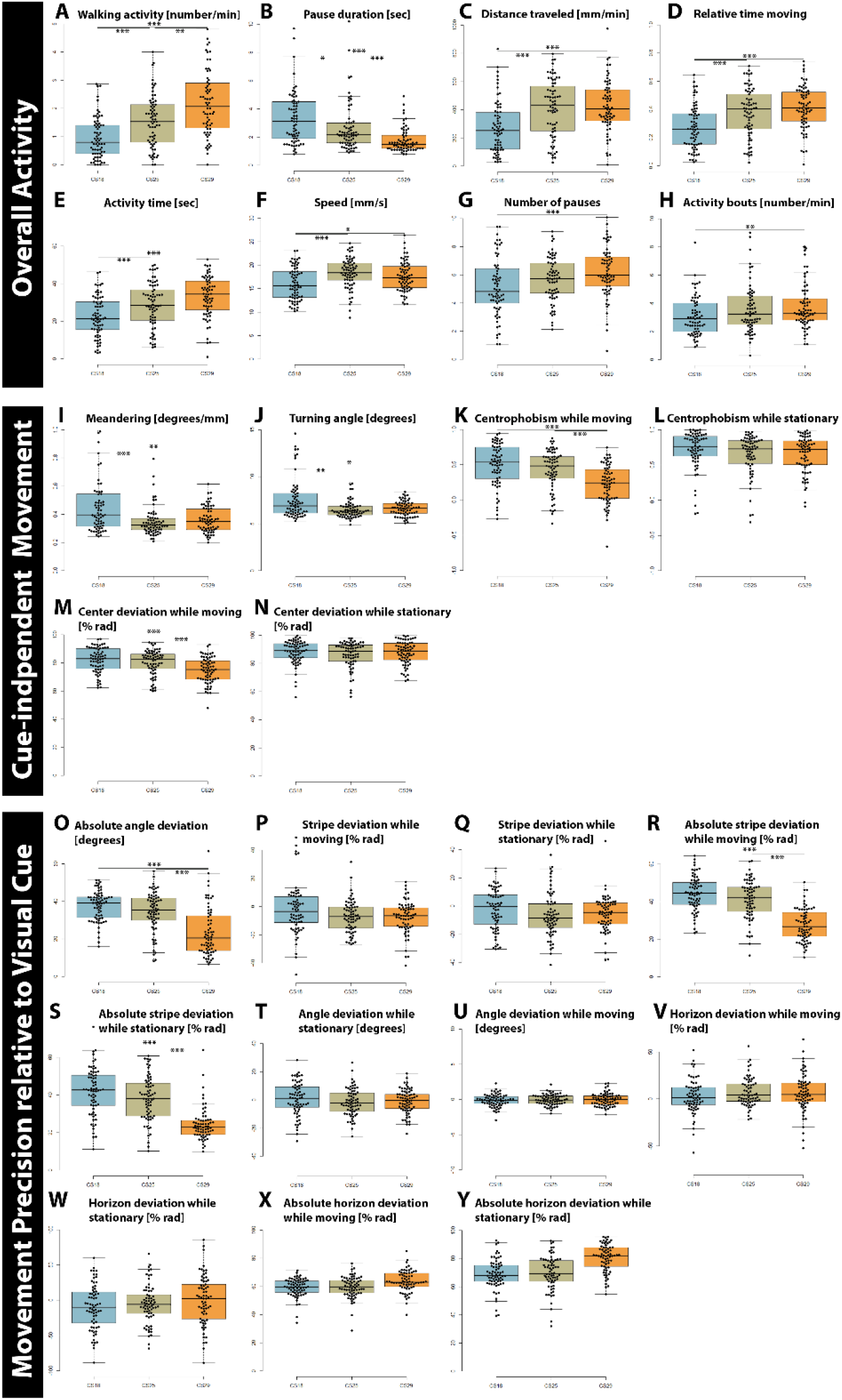
Analysis of 25 behavioral parameters in Buridan’s paradigm for temperature robustness. **(A-H)** Most parameters related to overall activity of flies in Buridan arena increase after development at high temperatures. **(I-Y)** Parameters related to cue-dependent and cue-independent movement precision are largely temperature-compensated. See Table S3 for quantifications.

**Table S1.** Summary of quantitative analyses of temperature effects in wild type

|  | 18 sem |  | 25 sem |  | 29 sem |  | 25/18 | 29/18 | 29/25 |
| --- | --- | --- | --- | --- | --- | --- | --- | --- | --- |
| R7 Filopodia speed (um/min) | **0,827** | 0,058 | **1,155** | 0,072 | **1,472** | 0,079 | **1,396614** | 1,779927 | 1,274459 |
| R7 Filopodia Lifetime (min) | **10,088** | 0,807 | **6,638** | 0,627 | **4,3** | 0,429 | **1,519735** | 2,346047 | 1,543721 |
| R7 BrpD3 active zones (#/terminal) | **24,65** | 0,523 | **20,65** | 0,323 | **16,35** | 0,597 | **1,193705** | 1,507645 | 1,262997 |
| R7 trans-Tango (# cell/optic lobe) | **183,7** | 7,106 | **145,3** | 7,027 | **133,8** | 6,335 | **1,264281** | 1,372945 | 1,085949 |
| R7-Dm8 GRASP (mean signal int.) | **0,17** | 0,008 | **0,137** | 0,006 | **0,116** | 0,005 | **1,240876** | 1,465517 | 1,181034 |
| R7-Dm9 GRASP (mean signal int.) | **0,093** | 0,009 | **0,066** | 0,006 | n/a | n/a | **1,409091** | n/a | n/a |
| R7-Dm11 GRASP (mean signal int.) | **0,115** | 0,011543 | **0,079** | 0,003 | n/a | n/a | **1,455696** | n/a | n/a |
| DCN Filopodia speed (um/min) | **0,235738736** | 0,01946 | **0,372921847** | 0,031406 | **0,761213** | 0,105939 | **1,581929** | 3,229053 | 2,041213 |
| DCN primary branch number | **7,074074074** | 0,417336 | **5,227272727** | 0,40649 | **2,62069** | 0,209537 | **1,353301** | 2,699318 | 1,994617 |
| DCN secondary branch number | **3,676470588** | 0,54257 | **1,484848485** | 0,267191 | **0,489796** | 0,115911 | **2,47599** | 7,506127 | 3,031566 |
| DCN BrpD3 active zones (#/terminal) | **0,275** | 0,008 | **0,241** | 0,009 | **0,104** | 0,01 | **1,141079** | 2,644231 | 2,317308 |
| DCN trans-Tango (# cells/optic lobe) | **88,83** | 9,964 | **55** | 2,637 | **13** | 2,65 | **1,615091** | 6,833077 | 4,230769 |
| DCN-Lawf1 GRASP (mean signal int.) | **0,259354806** | 0,009516 | **0,173174065** | 0,00545 | n/a | n/a | **1,497654** | n/a | n/a |
| R1-6 BrpD3 active zones (#/terminal) | **39,223** | 0,595 | **34,097** | 0,668 | **30,548** | 0,637 | **1,150336** | 1,283979 | 1,116178 |
| R1-6 ERG on transient (mV) | **2,85** | 0,118433 | **2,205** | 0,114817 | **1,985** | 0,118383 | **1,292517** | 1,435768 | 1,110831 |
| R1-6 ERG depolarization (mV) | **8,855** | 0,283445 | **9,095** | 0,356996 | **9,46** | 0,273515 | **0,973612** | 0,936047 | 0,961416 |
| Dm8 branch numbers | **321,28** | 14,4 | **259,94** | 11,29 | **215,87** | 8,35 | **1,235978** | 1,488303 | 1,204151 |
| Dm8 total branch length | **505,5074801** | 25,38734 | **429,0747891** | 20,30404 | **328,7044** | 16,81363 | **1,178134** | 1,537878 | 1,305351 |
| Dm8 overlap with #R7s | **14,73333333** | 0,693278 | **11,76470588** | 0,525311 | **10,23529** | 0,565747 | **1,252333** | 1,439464 | 1,149425 |
| Mi4 branch numbers | **284,12** | 9,06 | **246,43** | 8,93 | **197** | 6,14 | **1,152944** | 1,442234 | 1,250914 |
| Mi4 total branch length | **397,4117872** | 12,90929 | **349,6517454** | 11,07318 | **274,7308** | 7,369114 | **1,136593** | 1,44655 | 1,272707 |

**Table S2.** Summary of quantitative analyses of temperature effects in DIPγ mutants

|  | **DIPy cont** | **sem** | **DIPy null** | **sem** | **DIPy null/control** |
| --- | --- | --- | --- | --- | --- |
| R7-Dm8 GI | **0,128667** | 0,010655 | **0,07499** | 0,007825 | **0,58282438** |
| R7-Dm9 GI | **0,07957** | 0,005975 | **0,122925** | 0,00872 | **1,544865632** |
| R7-Dm11 C | **0,087512** | 0,005743 | **0,147891** | 0,012554 | **1,689949474** |
| R7-Mi1 GR | **0,008178** | 0,000401 | **0,080417** | 0,005984 | **9,833502068** |
| R7-Mi4 GR | **0,007972** | 0,000409 | **0,045601** | 0,003249 | **5,719861385** |
| R7-Tm9 GF | **0,006906** | 0,000597 | **0,041563** | 0,002742 | **6,018475543** |

**Table S3.**
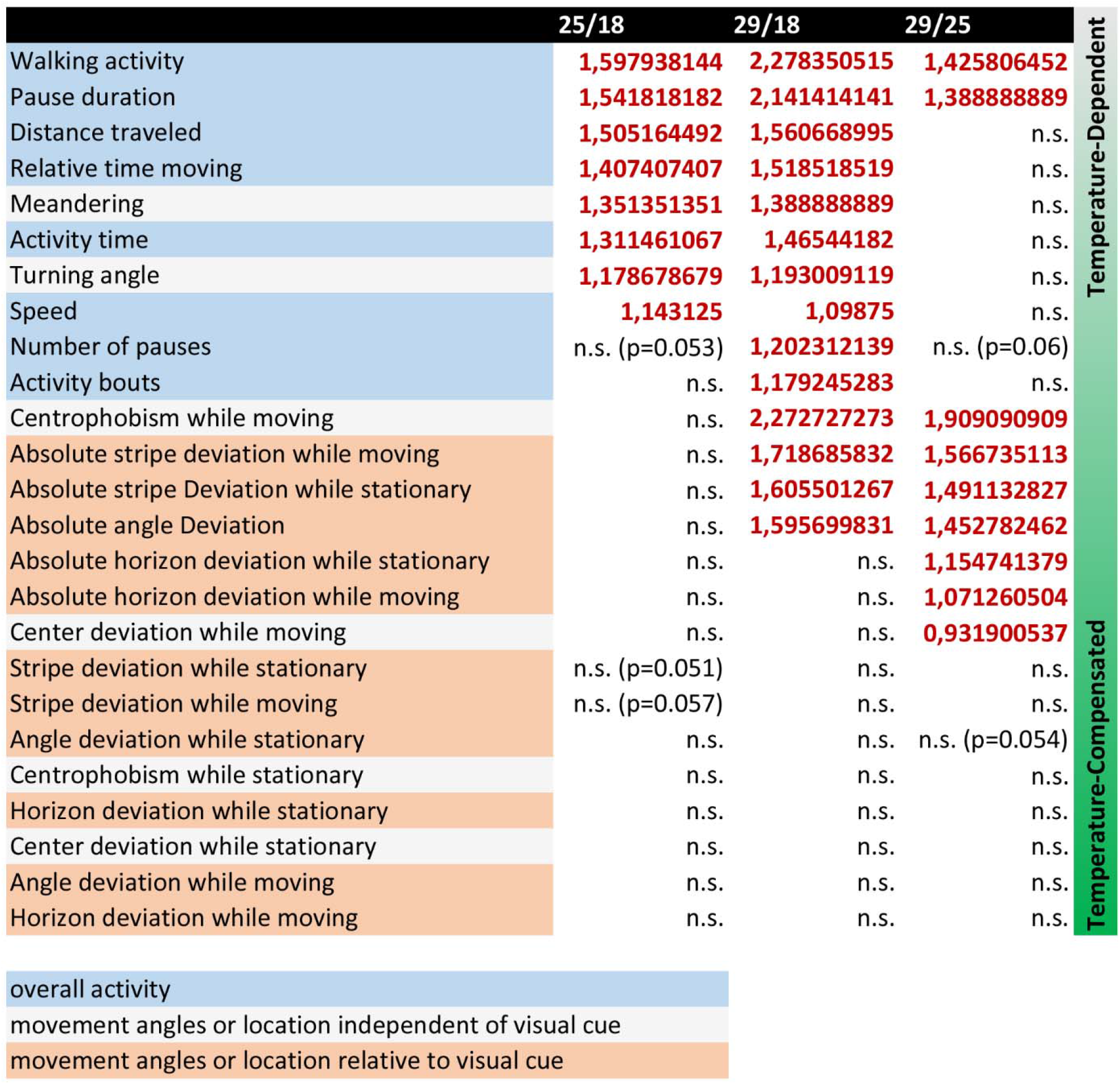
Summary of quantitative analyses of behavioral parameters

**Movie S1**. R7 axon filopodial dynamics at different temperatures

**Movie S2**. DCN axon branch dynamics at different temperatures

